# The Population Genetics of Collateral Resistance and Sensitivity

**DOI:** 10.1101/2020.08.25.267484

**Authors:** Sarah M. Ardell, Sergey Kryazhimskiy

## Abstract

Resistance mutations against one drug can elicit collateral sensitivity against other drugs. Multi-drug treatments exploiting such trade-offs can help slow down the evolution of resistance. However, if mutations with diverse collateral effects are available, a treated population may evolve either collateral sensitivity or collateral resistance. How to design treatments robust to such uncertainty is unclear. We show that many resistance mutations in *Escherichia coli* against various antibiotics indeed have diverse collateral effects. We propose to characterize such diversity with a joint distribution of fitness effects (JDFE) and develop a theory for describing and predicting collateral evolution based on simple statistics of the JDFE. We show how to robustly rank drug pairs to minimize the risk of collateral resistance and how to estimate JDFEs. In addition to practical applications, these results have implications for our understanding of evolution in variable environments.

## Introduction

The spread of resistance against most antibiotics and the difficulties in developing new ones has sparked considerable interest in using drug combinations and sequential drug treatments (Pál et al., 2015). Treatments where the drugs are chosen so that resistance against one of them causes the pathogen or cancer population to become sensitive to the other—a phenomenon known as collateral sensitivity—can eliminate the population before multi-drug resistance emerges (Pál et al., 2015; Pluchino et al., 2012).

Successful multi-drug treatments hinge on knowing which drugs select for collateral sensitivity against which other drugs. This information is obtained empirically by exposing bacterial and cancer-cell populations to drugs and observing the evolutionary outcomes (Bergstrom et al., 2004; Roemhild et al., 2020; Jensen et al., 1997; Imamovic and Sommer, 2013; Lázár et al., 2018; Maltas and Wood, 2019;Batra et al., 2021). Unfortunately, different experiments often produce collateral sensitivity profiles that are inconsistent with each other (e.g.,Imamovic and Sommer, 2013;Oz et al., 2014;Barbosa et al., 2017; Maltas and Wood, 2019). Some inconsistencies can be attributed to the fact that resistance mutations vary between bacterial strains, drug dosages, etc. (Mira et al., 2015; Barbosa et al., 2017; Das et al., 2020; Pinheiro et al., 2021; Card et al., 2020; Gjini and Wood, 2021). However, wide variation in collateral outcomes is observed even between replicate populations (Oz et al., 2014; Barbosa et al., 2017; Maltas and Wood, 2019; Nichol et al., 2019). This variation suggests that bacteria and cancers have access to multiple resistance mutations with different collateral sensitivity profiles, such that replicates can accumulate different mutations simply due to the intrinsic randomness of the evolutionary process (Jerison et al., 2020). However, the variability of collateral effects among resistance mutations has not been characterized (but see Card et al., 2021), and there is no principled approach for accounting for this variability in designing robust multi-drug treatments. In particular, it is unclear which evolutionary parameters determine the expected collateral outcomes of evolution and, importantly, the uncertainty around these expectations.

To address this problem, here we develop a population genetics theory of evolution of collateral sensitivity and resistance. Collateral sensitivity and resistance are specific examples of the more general evolutionary phenomenon, pleiotropy, which refers to any situation when one mutation affects multiple phenotypes (Wagner and Zhang, 2011; Paaby and Rockman, 2013). In case of drug resistance evolution, the direct effect of resistance mutations is to increase fitness in the presence of one drug (the “home” environment). In addition, they may also provide pleiotropic gains or losses in fitness in the presence of other drugs (the “non-home” environments) leading to collateral resistance or sensitivity, respectively.

Classical theoretical work on pleiotropy has been done in the field of quantitative genetics (Lande and Arnold, 1983; Rose, 1982; Barton, 1990; Slatkin and Frank, 1990; Jones et al., 2003; Johnson and Barton, 2005). In these models, primarily developed to understand how polygenic traits respond to selection in sexual populations, pleiotropy manifests itself as a correlated temporal change in multiple traits in a given environment. The question of how new strongly beneficial mutations accumulating in one environment affect the fitness of an asexual population in future environments is outside of the scope of these models. The pleiotropic consequences of adaptation have also been explored in various “fitness landscape” models (e.g. Connallon and Clark, 2015; Martin and Lenormand, 2015; Harmand et al., 2017; Wang and Dai, 2019; Maltas et al., 2019; Tikhonov et al., 2020). This approach helps us understand how evolutionary trajectories and outcomes depend on the global structure of the underlying fitness landscape. However, it is difficult to use these models to predict collateral outcomes because the global structure of fitness landscape is unknown and notoriously difficult to estimate even in controlled laboratory conditions.

Here, we take a different approach which is agnostic with respect to the global structure of the fitness landscape. Instead, we assume only the knowledge of the so-called joint distribution of fitness effects (JDFE), i.e., the probability that a new mutation has a certain pair of fitness effects in the home and non-home environments (Jerison et al., 2014; Martin and Lenormand, 2015; Bono et al., 2017). JDFE is a natural extension of the DFE, the distribution of fitness effects of new mutations, often used in modeling evolution in a single environment (King, 1972; Ohta, 1987; Orr, 2003; Rees and Bataillon, 2006; Eyre-Walker and Keightley, 2007; Martin and Lenormand, 2008; MacLean and Buckling, 2009; Kryazhimskiy et al., 2009; Levy et al., 2015). Like the DFE, the JDFE is a local property of the fitness landscape which means that it can be at least in principle estimated, for example using a variety of modern high-throughput techniques (e.g.,Qian et al., 2012; Hietpas et al., 2013; Van Opijnen et al., 2009; Stiffler et al., 2015; Chevereau et al., 2015; Levy et al., 2015; Blundell et al., 2019; Bakerlee et al., 2021). The downside of this approach is that the JDFE can change over time as the population traverses the fitness landscape (Good et al., 2017; Venkataram et al., 2020; Aggeli et al., 2020). However, in the context of collateral drug resistance and sensitivity, we are primarily interested in short time scales over which JDFE can be reasonably expected to stay approximately constant.

The rest of the paper is structured as follows. First, we use previously published data to demonstrate that the bacterium *Escherichia coli* has access to drug resistance mutations with diverse collateral effects. This implies that, rather than treating collateral effects as deterministic properties of drug pairs, we should think of them probabilistically, in terms of the respective JDFEs. We then show that a naive intuition about how the JDFE determines pleiotropic outcomes of evolution can sometimes fail, and a rigorous mathematical approach is therefore required. We develop such an approach, which reveals two key “pleiotropy statistics” of the JDFE that determine the dynamics of fitness in the non-home condition. Our theory makes quantitative predictions in a variety of regimes if the population genetic parameters are known. However, we argue that in the case of drug resistance evolution the more important problem is to robustly order drug pairs in terms of their collateral sensitivity profiles even if the population genetic parameters are unknown.

We develop a metric that allows us to do so. Finally, we provide some practical guidance for estimating the pleiotropy statistics of empirical JDFEs in the context of ranking drug pairs.

## Results

### Antibiotic resistance mutations in *E. coli* have diverse collateral effects

We begin by demonstrating that JDFE is a useful concept for modeling the evolution of collateral antibiotic resistance and sensitivity. If all resistance mutations against a given drug had identical pleiotropic effects on the fitness of the organism in presence of another drug, the dynamics of collateral resistance/sensitivity could be understood without the JDFE concept. On the other hand, if different resistance mutations have different pleiotropic fitness effects, predicting the collateral resistance/sensitivity dynamics requires specifying the probabilities with which mutations with various home and non-home fitness effects arise in the population. The JDFE specifies these probabilities. Therefore, for the JDFE concept to be useful in the context of collateral resistance/sensitivity evolution, we need to show that resistance mutations against common drugs have diverse collateral effects in the presence of other drugs.

To our knowledge, no data sets are currently publicly available that would allow us to systematically explore the diversity of collateral effects among all resistance mutations against any one drug in any organism. Instead, we examined the fitness effects of 3883 gene knock-out mutations in the bacterium *Escherichia coli*, measured in the presence of six antibiotics (Chevereau et al., 2015), as well as the fitness effects of 4997 point mutations in the TEM-1 *β*-lactamase gene measured in the presence of two antibiotics (Stiffler et al, 2015).

For the four out of six antibiotics used by Chevereau et al. (2015), we find between 12 (0.31 %) and 170 (4.38 %) knock-out mutations that provide some level of resistance against at least one of the antibiotics (false discovery rate (FDR) ~ 25%; Figure 1, Supplementary Table S1; see Materials and Methods for details). Plotting on the *x*-axis the fitness effect of each knock-out mutation in the presence of the drug assumed to be applied first (i.e., the home environment) against its effect in the presence of another drug assumed to be applied later (i.e., the non-home environment, *y*-axis), we find mutations in all four quadrants of this plane, for all 12 ordered drug pairs (Figure 1, Supplementary Table S1). Similarly, we find diverse collateral effects among mutations within a single gene (Figure S1; see Materials and Methods for details).

**Figure 1.**
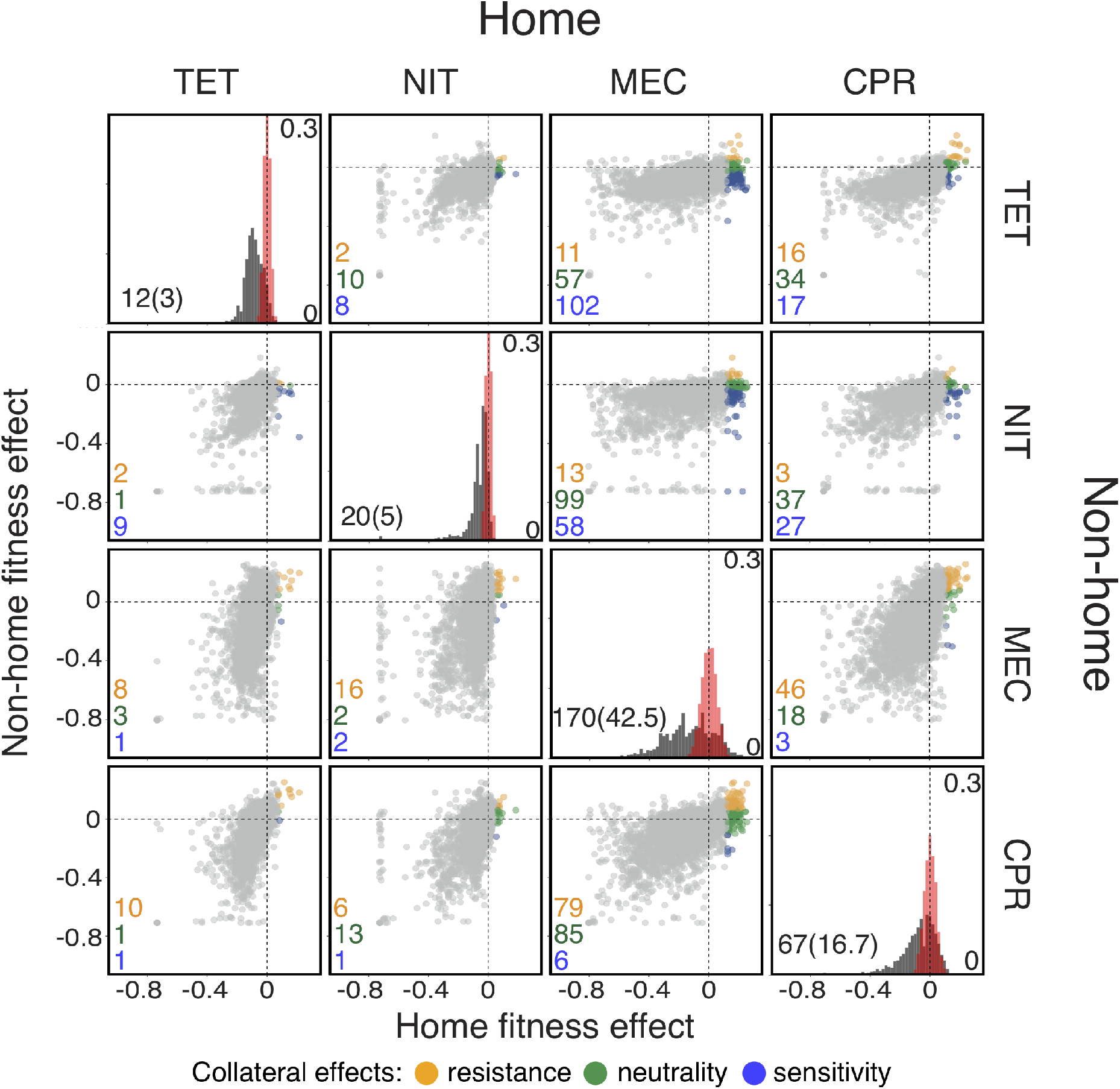
Fitness effects of gene knock-out mutations in *E. coli* in the presence of four antibiotics. Data are from Chevereau et al. (2015). Each diagonal panel shows the distribution of fitness effects (DFE) of knock-out mutations in the presence of the corresponding antibiotic (equivalent to Figure 1C in Chevereau et al. (2015)). Scale of the *y*-axis in these panels is indicated inside on the right. The estimated measurement noise distributions are shown in red (see Materials and Methods for details). Note that some noise distributions are vertically cut-off for visual convenience. The number of identified beneficial mutations (i.e., resistance mutations) and the expected number of false positives (in parenthesis) are shown in the bottom left corner. The list of identified resistance mutations is given in the Supplementary Table S1. Off-diagonal panels show the fitness effects of knock-out mutations across pairs of drug environments. The *x*-axis shows the fitness in the environment where selection would happen first (i.e., the “home” environment). Each point corresponds to an individual knock-out mutation. Resistance mutations identified in the home environment are colored according to their collateral effects, as indicated in the legend. The numbers of mutations of each type are shown in the corresponding colors in the bottom left corner of each panel. TET: tetracycline; NIT: nitrofurantoin; MEC: mecillinam; CPR: ciprofloxacin.

Since both data sets represent subsets of all resistance mutations, we conclude that *E. coli* likely have access to resistance mutations with diverse pleiotropic effects, such that a fitness gain in the presence of any one drug can come either with a pleiotropic gain or a pleiotropic loss of fitness in the presence of other drugs. Therefore, the JDFE framework is suitable for modeling the evolution of collateral resistance/sensitivity. In the next section, we formally define a JDFE and probe our intuition for how its shape determines the fitness trajectories in the non-home environment.

### JDFE determines the pleiotropic outcomes of adaptation

For any genotype *g* that finds itself in one (“home”) environment and may in the future encounter another “non-home” environment, we define the JDFE as the probability density Φ_g_(Δ*x*, Δ*y*) that a new mutation that arises in this genotype has the selection coefficient Δ*x* in the home environment and the selection coefficient Δ*y* in the non-home environment (Jerison et al., 2014). For concreteness, we define the fitness of a genotype as its malthusian parameter (Crow and Kimura, 1972). So, if the home and non-home fitness of genotype *g* are *x* and *y*, respectively, and if this genotype acquires a mutation with selection coefficients Δ*x* and Δ*y*, its fitness becomes *x* + Δ*x* and *y* + Δ*y*. This definition of the JDFE can, of course, be naturally extended to multiple non-home environments. In principle, the JDFE can vary from one genotype to another. However, to develop a basic intuition for how the JDFE determines pleiotropic outcomes, we assume that all genotypes have the same JDFE. We discuss possible extensions to epistatic JDFEs in Appendix A.

The JDFE is a complex object. So, we first ask whether some simple and intuitive summary statistics of the JDFE may be sufficient to predict the dynamics of the non-home fitness of a population which is adapting in the home environment. Intuitively, if there is a trade-off between home and non-home fitness, non-home fitness should decline; if the opposite is true, non-home fitness should increase. Canonically, a trade-off occurs when any mutation that improves fitness in one environment decreases it in the other environment and vice versa (Roff and Fairbairn, 2007). Genotypes that experience such “hard” trade-offs are at the Pareto front (Shoval et al., 2012; Li et al., 2019). For genotypes that are not at the Pareto front, some mutations that are beneficial in the home environment may be beneficial in the non-home environment and others may be deleterious. In this more general case, trade-offs are commonly quantified by the degree of negative correlation between the effects of mutations on fitness in the two environments (Roff and Fairbairn, 2007; Tikhonov et al., 2020). Thus, we might expect that evolution on negatively correlated JDFEs would lead to pleiotropic fitness losses and evolution on positively correlated JDFEs would lead to pleiotropic fitness gains.

To test this intuition, we generated a family of Gaussian JDFEs that varied, among other things, by their correlation structure (Figure 2; Materials and Methods). We then simulated the evolution of an asexual population on these JDFEs using a standard Wright-Fisher model (Materials and Methods) and tested whether the trade-off strength, measured by the JDFE’s correlation coefficient, predicts the dynamics of non-home fitness. Figure 2 shows that our naive expectation is incorrect. Positively correlated JDFEs sometimes lead to pleiotropic fitness losses (Figure 1D, I), and negatively correlated JD-FEs sometimes lead to pleiotropic fitness gains (Figure 2B,G). Even if we calculate the correlation coefficient only among mutations that are beneficial in the home environment, the pleiotropic outcomes still do not always conform to the naive expectation, as the sign of the correlation remains the same as for the full JDFEs in all these examples.

**Figure 2.**
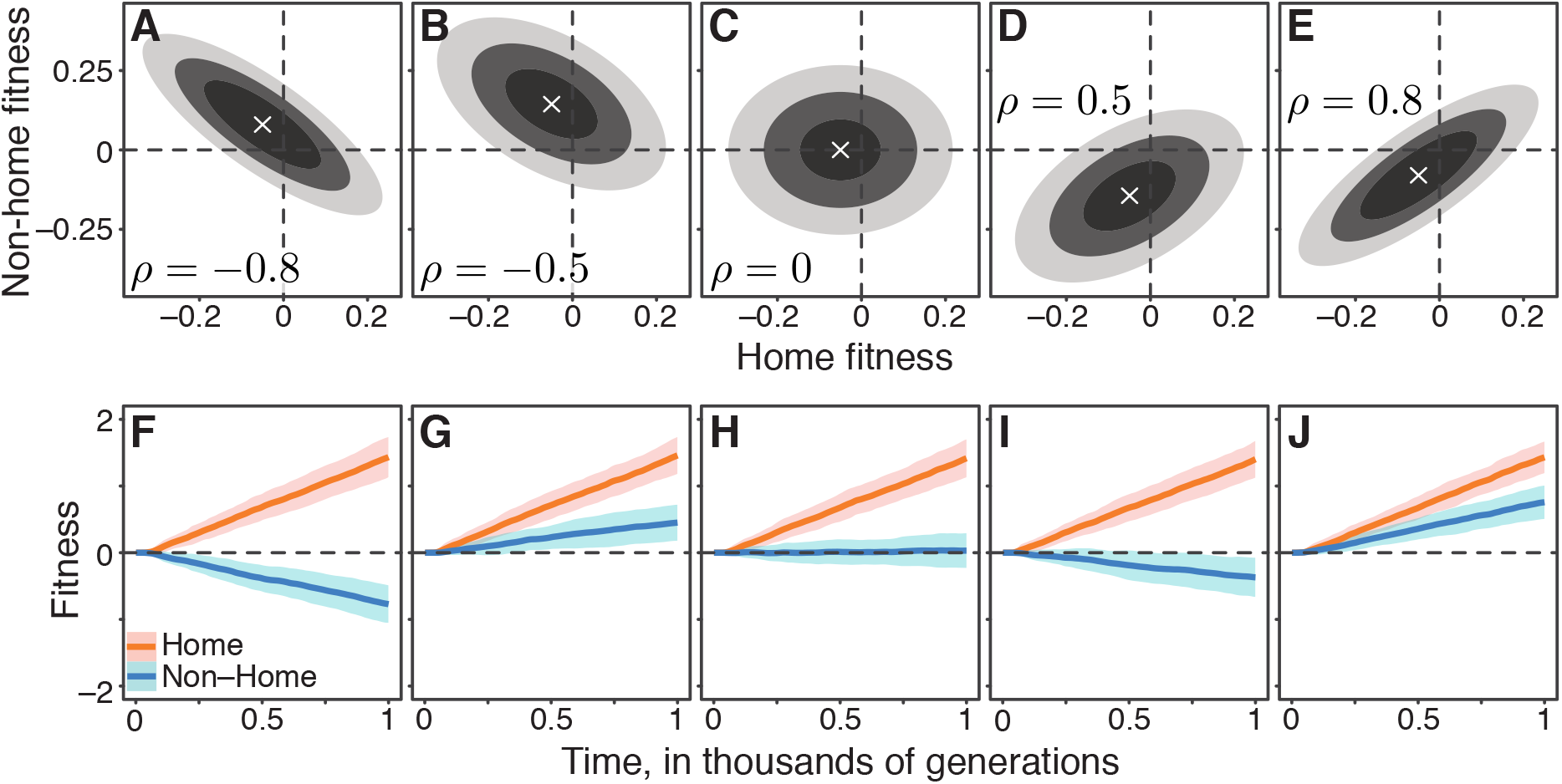
Gaussian JDFEs and the resulting fitness trajectories. **A**–**E.** Contour lines for five Gaussian JDFEs. “x” marks the mean. For all distributions, the standard deviation is 0.1 in both home- and non-home environments. The correlation coefficient *ρ* is shown in each panel. **F**–**J.** Home and non-home fitness trajectories for the JDFEs shown in the corresponding panels above. Thick lines show the mean, ribbons show ±1 standard deviation estimated from 100 replicate simulations. Population size *N* = 10^4^, mutation rate *U* = 10^−4^ (*U_b_* = 4.6 ×10^−5^).

There are other properties of the JDFE that we might intuitively expect to be predictive of the pleiotropic outcomes of adaptation. For example, among the JDFEs considered in Figure 2, it is apparent that those with similar relative probability weights in the first and fourth quadrants produce similar pleiotropic outcomes. However, simulations with other JDFE shapes show that even distributions that are similar according to this metric can also result in qualitatively different pleiotropic outcomes (Supplementary Figure S2).

Overall, our simulations show that JDFEs with apparently similar shapes can produce qualitatively different trajectories of pleiotropic fitness changes (e.g., compare Figures 2A,F and 2B,G or Figures 2D,I and 2E,J). Conversely, JDFEs with apparently different shapes can result in rather similar pleiotropic outcomes (e.g., compare Figures 2B,G and 2E,J or Figures 2A,F and 2D,I). Thus, while the overall shape of the JDFE clearly determines the trajectory of pleiotropic fitness changes, it is not immediately obvious what features of its shape play the most important role, particularly if the JDFE is more complex than a multivariate Gaussian. In other words, even if we have perfect knowledge of the fitness effects of all mutations in multiple environments, converting this knowledge into a qualitative prediction of the expected direction of pleiotropic fitness change (gain or loss) does not appear straightforward. Therefore, we next turn to developing a population genetics model that would allow us to predict not only the direction of pleiotropic fitness change but also the expected rate of this change and the uncertainty around the expectation.

### The population genetics of pleiotropy

To systematically investigate which properties of the JDFE determine the pleiotropic fitness changes in the non-home environment, we consider a population of size *N* that evolves on a JDFE in the “strong selection weak mutation” (SSWM) regime, also known as the “successional mutation” regime (Orr, 2000; Desai and Fisher, 2007; Kryazhimskiy et al, 2009; Good and Desai, 2015).

We consider an arbitrary JDFE without epistasis, that is a situation when all genotypes have the same JDFE Φ (Δ*x*, Δ*y*). We explore an extension to JDFEs with simple forms of epistasis in Appendix A. We assume that mutations arise at rate *U* per individual per generation. In the SSWM limit, a mutation that arises in the population either instantaneously fixes or instantaneously dies out. Therefore, the population is essentially monomorphic at all times, such that at any time *t* we can characterize it by its current pair of fitness values (*X_t_,Y_t_*). If a new mutation with a pair of selection coefficients (Δ*x*, Δ*y*) arises in the population at time *t*, it fixes with probability 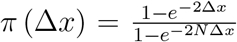 (Kimura, 1962) in which case the population’s fitness transitions to a new pair of values
(*X_t_* + Δ*x*, *Y_t_* + Δ*y*). If the mutation dies out, an event that occurs with probability 1 – π (Δ*x*), the population’s fitness does not change. This model specifies a continuous-time two-dimensional Markov process.

In general, the dynamics of the probability density *p*(*x,y,t*) of observing the random vector (*X_t_*, *Y_t_*) at values (*x, y*) are governed by an integro-differential forward Kolmogorov equation, which is difficult to solve (Materials and Methods). However, if most mutations that contribute to adaptation have small effects, these dynamics are well approximated by a diffusion equation which can be solved exactly (Materials and Methods). Then *p*(*x, y, t*) is a normal distribution with mean vector

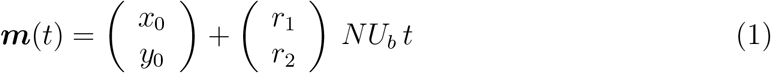

and variance-covariance matrix

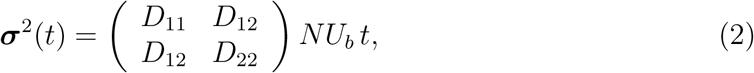

where are *r*_1_ and *r*_2_, given by equations (7) and (8) in Materials and Methods, are the expected fitness effects in the home and non-home environments for a mutation fixed in the home environment, and *D*_11_, *D*_12_ and *D*_22_, given by equations (9)–(11) in Materials and Methods, are the second moments of this distribution. Here, 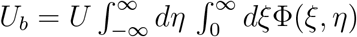 is the total rate of mutations beneficial in the home environment, and *x*_0_ and *y*_0_ are the initial values of population’s fitness in the home and non-home environments.

Equations (1), (2) show that the distribution of population’s fitness at time *t* in the non-home environment is entirely determined by two parameters, *r*_2_ and *D*_22_, which we call the pleiotropy statistics of the JDFE. The expected rate of fitness change in the non-home environment depends on the pleiotropy statistic *r*_2_, which we refer to as the expected pleiotropic effect. Thus, evolution on a JDFE with a positive *r*_2_ is expected to result in pleiotropic fitness gains and evolution on a JDFE with a negative *r*_2_ is expected to result in pleiotropic fitness losses. Equation (2) shows that the variance around this expectation is determined by the pleiotropy variance statistic *D*_22_. Since both the expectation and the variance change linearly with time (provided *r*_2_ ≠ 0), the change in the non-home fitness in any replicate population would eventually have the same sign as *r*_2_, but the time scale of such convergence depends on the “collateral risk” statistic 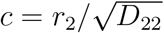 (Materials and Methods). This observation has important practical implications, and we return to it in the Section “Robust ranking of drug pairs”.

These theoretical results suggest a simple explanation for the somewhat counter-intuitive observations in Figure 2. We may intuitively believe that evolution on negatively correlated JDFEs should lead to fitness losses in the non-home environment because on such JDFEs mutations with largest fitness benefits in the home environment typically have negative pleiotropic effects. However, such mutations may be too rare to drive adaptation. At the same time, the more common mutations that do typically drive adaptation may have positive pleiotropic effects, in which case the population would on average gain non-home fitness, as in Figure 2B. Our theory shows that to predict the direction of non-home fitness change, the frequency of beneficial mutations with various pleiotropic effects and the strength of these effects need to be weighted by the likelihood that these mutations fix. The expected pleiotropic effect *r*_2_ accomplishes this weighting.

We tested the validity of equations (1) and (2) by simulating evolution in the SSWM regime on 125 Gaussian JDFEs with various parameters (Materials and Methods) and found excellent agreement (Figure 3A,B). However, many microbes likely evolve in the “concurrent mutation” regime, i.e., when multiple beneficial mutations segregate in the population simultaneously (Desai and Fisher, 2007; Lang et al., 2013). As expected, our theory fails to quantitatively predict the pleiotropic fitness trajectories when *NU_b_* > 1 (Supplementary Figure S3). However, the expected rate of change of non-home fitness and its variances remain surprisingly well correlated with the pleiotropy statistics *r*_2_ and *D*_22_ across various JDFEs (Supplementary Figure S3). In other words, we can still use these statistics to correctly predict whether a population would lose or gain fitness in the non-home environment and to order the non-home environments according to their expected pleiotropic fitness changes and variances. We will exploit the utility of such ranking in the next section.

**Figure 3.**
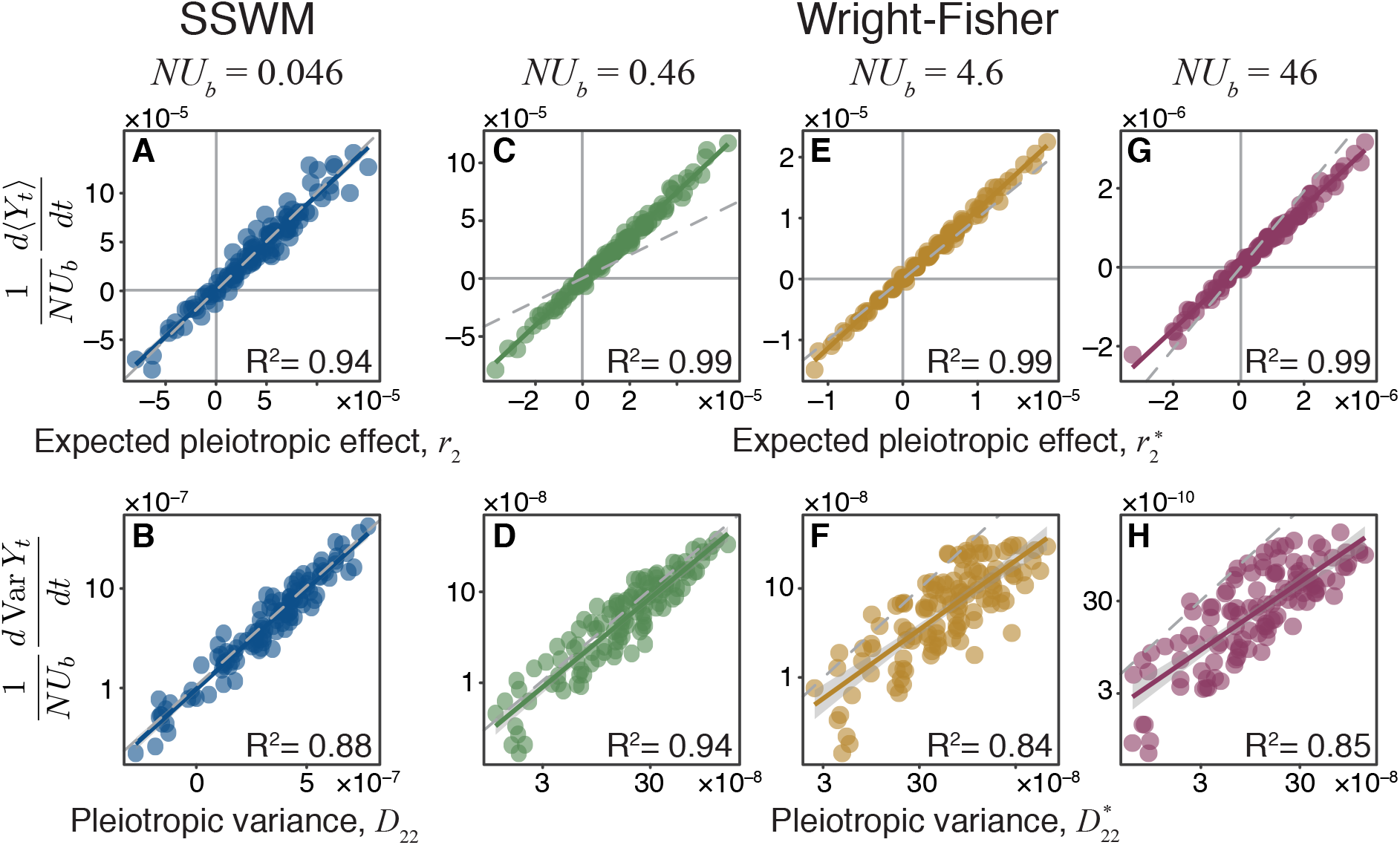
Pleiotropy statistics predict the properties of non-home fitness trajectories in simulations. Each point corresponds to an ensemble of replicate simulation runs with the same population genetic parameters on one of 125 Gaussian JDFEs (see Supplementary Table S3 for the JDFE parameters). **A.** Expected pleiotropic effect *r*_2_ versus the scaled slope of the mean rate of non-home fitness change observed in SSWM simulations. **B.** Pleiotropic variance *D*_22_ versus the scaled rate of change in the variance in non-home fitness observed in SSWM simulations. **C, E, G.** Expected pleiotropic effect 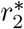 versus the scaled slope of the mean rate of non-home fitness change observed in Wright-Fisher simulations. **D, F, H.** Pleiotropic variance 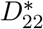 versus the scaled rate of change in the variance in non-home fitness observed in Wright-Fisher simulations simulations. (See Supplementary Figure S3 for comparison between simulations and the unadjusted pleiotropy statistics *r*_2_ and *D*_22_.) 1000 replicate simulations were carried out in the SSWM regime. All Wright-Fisher simulations were carried out with *U* = 10^−4^ and variable *N*, 300 replicate simulations per data point. (see Materials and Methods for details). In all panels, the grey dashed line represents the identity (slope 1) line, and the solid line of the same color as the points is the linear regression for the displayed points (*R*^2^ value is shown in each panel; *P* < 2 ×10^−16^ for all regressions).

We next sought to expand our theory to the concurrent mutation regime. A key characteristic of adaptation in this regime is that mutations whose fitness benefits in the home environment are below a certain “effective neutrality” threshold are usually outcompeted by superior mutations and therefore fix with lower probabilities than predicted by Kimura’s formula (Schiffels et al., 2011; Good et al., 2012). Good et al. (2012) provide an equation for calculating the fixation probability π* (Δ*x*) for a mutation with home fitness benefit Δ*x* in the concurrent mutation regime (equation (6) in Good et al. (2012)). Thus, by replacing 2*ξ* (the approximate fixation probability in the SSWM regime) in equations (8) and (11) with π*(*ξ*), we obtain the adjusted pleiotropy statistics 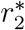 and 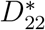 for the concurrent mutation regime (see Materials and Methods for details).

To test how well these statistics predict the dynamics of fitness in the non-home environment, we simulated evolution on the same 125 JDFEs using the full Wright-Fisher model with a range of population genetic parameters that span the transition from the successional to the concurrent mutation regimes for 1000 generations. We find that 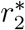 quantitatively predicts the expected rate of non-home fitness change, with a similar accuracy as Good et al. (2012) predict the rate of fitness change in the home environment, as long as *NU_b_* > 1 (Figure 3C,E,G; compare with Figure S3A,C,E). 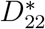 also predicts the empirically observed variance in non-home fitness trajectories much better than *D*_22_, although this relationship is more noisy than between mean fitness and 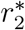 (Figure 3D,F,H; compare with Figure S3B,D,F). Some of this noise can be attributed to sampling, as we estimate both the mean and the variance from 300 replicate simulation runs, and the variance estimation is more noisy. Even in the absence of sampling noise however, we do not expect that 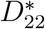 would predict the non-home fitness variance perfectly because our theory does not account for the autocorrelation in the fitness trajectories that arise in the concurrent mutation regime but not in the successive mutation regime (see Appendix D in Desai and Fisher (2007)). To our knowledge, the correct analytical calculation for fitness variance even in the home environment is not yet available.

Overall, our theory allows us to quantitatively predict the dynamics of non-home fitness in a range of evolutionary regimes if the JDFE and the population genetic parameters *N* and *U_b_* are known. However, neither the full JDFE nor the population genetic parameters will likely be known in most practical situations, such as designing a drug treatment for a cancer patient. In the next section, we address the question of how to robustly select drug pairs for a sequential treatment, assuming that the pleiotropy statistics *r*_2_ and *D*_22_ are known but the population genetic parameters are not. In the Section “Measuring JDFEs” we provide some guidance on how the JDFE can be measured.

### Robust ranking of drug pairs

Consider a hypothetical scenario where a drug treatment is being designed for a patient with a tumor or a bacterial infection. In selecting a drug, it is desireable to take into account not only the standard medical considerations, such as drug availability, toxicity, etc., but also the possibility that the treatment with this drug will fail due to the evolution of resistance. Therefore, it may be prudent to consider a list of drugs pairs (or higher-order combinations), ranked by the propensity of the first drug in the pair to elicit collateral resistance against the second drug in the pair. All else being equal, the drug deployed first should form a high-ranking pair with at least one other secondary drug. Then, if the treatment with the first drug fails, a second one can be deployed with a minimal risk of collateral resistance. Thus, we set out to develop a metric for ranking drug pairs according to this risk.

Clearly, any drug pair with a negative *r*_2_ is preferable over any drug pair with a positive *r*_2_, since the evolution in the presence of the first drug in a pair with *r*_2_ < 0 is expected to elicit collateral sensitivity against the second drug in the pair but the opposite is true for drug pairs with *r*_2_ > 0. It is also clear that among two drug pairs with negative *r*_2_, a pair with a more negative *r*_2_ and lower *D*_22_ is preferable over a pair with a less negative *r*_2_ and higher *D*_22_ because evolution in the presence of the first drug in the former pair will more reliably lead to stronger collateral sensitivity against the second drug in the pair. The difficulty is in how to compare and rank two drug pairs where one pair has a more negative *r*_2_ but higher *D*_22_. Our theory shows that the chance of emergence of collateral resistance monotonically increases with the collateral risk statistic 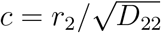 (see Materials and Methods). Thus, we propose to rank drug pairs by *c* from lowest (most negative and therefore most preferred) to highest (least negative or most positive and therefore least preferred).

To demonstrate the utility of such ranking, consider four hypothetical drug pairs with JDFEs shown in Figure 4A. The similarity between their shapes makes it difficult to predict *a priori* which one would have the lowest and highest probabilities of collateral resistance. Thus, we rank these JDFEs by their *c* statistic. To test whether this ranking is accurate with respect to the risk of collateral resistance, we simulate the evolution of a Wright-Fisher population in the presence of the first drug in each pair for 600 generations and estimate the probability that the evolved population has a positive fitness in the presence of the second drug, i.e., the probability that it becomes collaterally resistant (Figure 4B). We find that our a priori ranking corresponds perfectly to the ranking according to this probability, evidenced by the consistent higher collateral resistance risk for JDFEs with higher *c* over time (Figure 4B). Interestingly, the top ranked JDFE does not have the lowest expected pleiotropic effect *r*_2_. Nevertheless, the fact that the pleiotropic variance statistic *D*_22_ for this JDFE is small ensures that the risk of collateral resistance evolution is the lowest. This 1 to 1 rank correlation holds more broadly, for all 125 Gaussian JDFEs and all population genetic parameters considered in the previous section (Figure 4D). In other words, we can use the collateral risk statistic *c* to robustly rank drug pairs according to the risk of collateral resistance evolution, irrespectively of the population genetic parameters.

**Figure 4.**
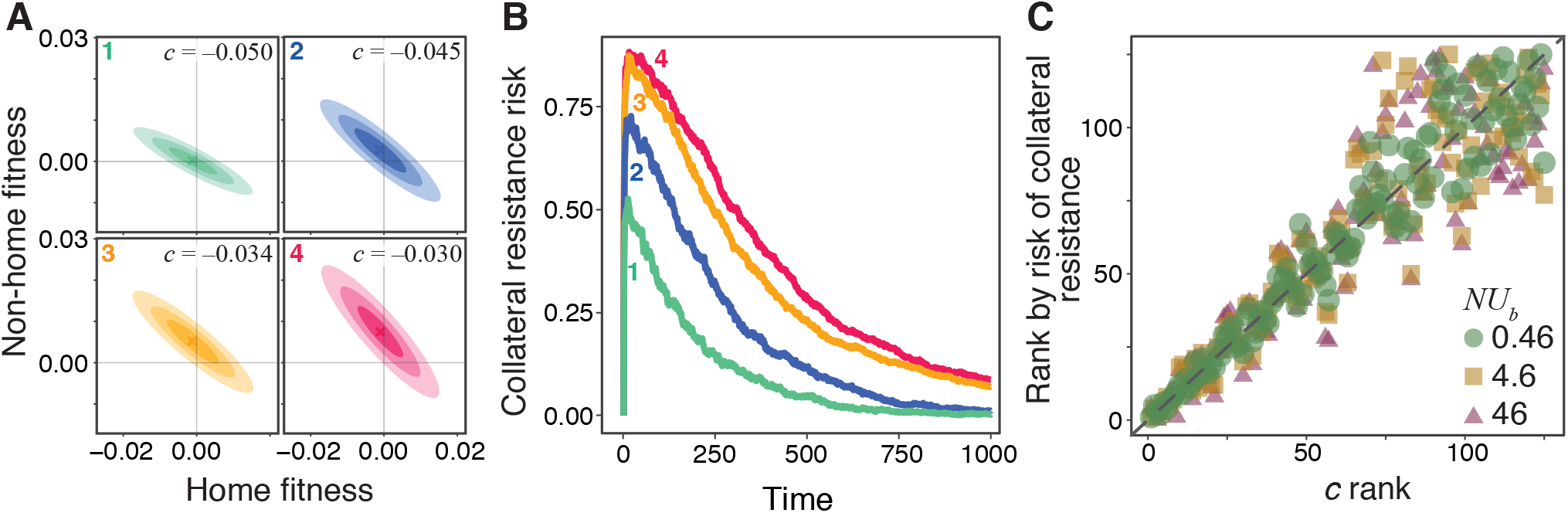
Robust ranking of drug pairs. **A.** Four hypothetical JDFEs, ranked by their ***c*** statistic. For all four JDFEs, the mean and the standard devion in the home environment are −1 × 10^−3^ and 0.01, respectively. The mean and the standard deviation in the non-home environment are 1 × 10^−4^ and 5.1 × 10^−3^ (rank 1), 2.6 × 10^−3^ and 7.5 × 10^−3^ (rank 2), 5.1 × 10^−3^ and 7.5 × 10^−3^ (rank 3), 7.5 × 10^−3^ and 0.01 (rank 4). Correlation coefficient for all four JDFEs is −0.9. **B.** Collateral resistance risk over time, measured as the fraction of populations with positive mean fitness in the non-home environment. These fractions are estimated from 1000 replicate Wright-Fisher simulation runs with *N* = 10^4^, *U* = 10^−4^ (*NU_b_* = 0.46). Colors correspond to the JDFEs in panel A. Numbers indicate the *c*-rank of each JDFE. **C.** *A priori c*-rank (*x*-axis) versus the *a posteriori* rank (*y*-axis) based on the risk of collateral resistance observed in simulations, for all 125 Gaussian JDFEs and all *NU_b_* values shown in Figure 3. Grey dashed line is the identity line.

### Measuring JDFEs

So far, we assumed that the parameters of the JDFE on which the population evolves are known. In reality, they have to be estimated from data, which opens up at least two practically important questions. The first question is experimental. From what types of data can JDFEs be in principle estimated and how good are different types of data for this purpose? We can imagine, for example, that some properties of JDFEs can be estimated from genome sequencing data (Jerison et al., 2020) or from temporally resolved fitness trajectories (Bakerlee et al., 2021). Here we focus on the most direct way of estimating JDFE parameters, from the measurements of the home and non-home fitness effects of individual mutations. The experimental challenge with this approach is to sample those mutations that will most likely contribute to adaptation in the home environment (see “Discussion” for an extended discussion of this problem). Below, we propose two potential strategies for such sampling: the Luria-Delbrück (LD) method and the barcode lineage tracking (BLT) method. The second question is statistical: how many mutants need to be sampled to reliably rank drug pairs according to the risk of collateral resistance? We evaluate both proposed methods with respect to this property.

The idea behind the LD method is to expose the population to a given drug at a concentration above the minimum inhibitory concentration (MIC), so that only resistant mutants survive (Pinheiro et al., 2021). This selection is usually done on agar plates, so that individual resistant mutants form colonies and can be isolated. The LD method is relatively easy to implement experimentally, but it is expected to work only if the drug concentration is high enough to kill almost all non-resistant cells. In reality, resistant mutants may be selected at concentrations much lower than MIC (Andersson and Hughes, 2014). Furthermore, mutants selected at different drug concentrations may be genetically and functionally distinct (Lindsey et al., 2013; Pinheiro et al., 2021) and therefore may have statistically different pleiotropic profiles. As a result, mutants sampled with the LD method may not be most relevant for predicting collateral evolution at low drug concentrations, and other sampling methods may be required for isolating weakly beneficial mutations.

Isolating individual weakly beneficial mutations is more difficult because by the time a mutant reaches a detectable frequency in the population it has accumulated multiple additional driver and passenger mutations (Lang et al., 2013; Nguyen Ba et al., 2019). One way to isolate many single beneficial mutations from experimental populations is by using the recently developed barcode lineage tracking (BLT) method (Levy et al., 2015; Venkataram et al., 2016). In a BLT experiment, each cell is initially tagged with a unique DNA barcode. As long as there is no recombination or other DNA exchange, any new mutation is permanently linked to one barcode. A new adaptive mutation causes the frequency of the linked barcode to grow, which can be detected by sequencing. By sampling many random mutants and genotyping them at the barcode locus, one can identify mutants from adapted lineages even if they are rare (Venkataram et al., 2016). As a result, BLT allows one to sample mutants soon after they acquire their first driver mutation, before acquiring secondary mutations.

To evaluate the quality of sampling based on the LD and BLT methods, we consider the following hypothetical experimental setup. *K* beneficial mutants are sampled from each home environment (with either one of the methods), and their home and non-home fitness (*X_i_, Y_i_*) are measured for each mutant *i* = 1,…, *K*. Since we are ultimately interested in ranking drug pairs by their risk of collateral resistance, we estimate the collateral risk statistic *ĉ* from these fitness data for each drug pair and use *ĉ* to rank them (see Materials and Methods for details). We compare such *a priori* ranking of 125 hypothetical drug pairs with Gaussian JDFEs used in previous sections with their *a posteriori* ranking based on the risk of collateral resistance observed in simulations.

To model the LD sampling method on a given JDFE, we randomly sample *K* mutants whose home fitness exceeds a certain cutoff. To model a BLT experiment, we simulate evolution in the home environment and randomly sample *K* beneficial mutants from generation 250 (see Materials and Methods for details). We find that the *ĉ*-ranking estimated with either LD or BLT methods captures the *a posteriori* ranking surprisingly well, even when the number of sampled mutants is as low as 10 per drug pair (Figure 5). Given that the JDFEs with adjacent ranks differ in c by a median of only 0.65%, the strong correlations shown in Figure 5 suggest that even very similar JDFEs can be differentiated with moderate sample sizes. As expected, this correlation is further improved upon increased sampling, and it is insensitive to the specific home fitness threshold that we use in the LD method (Figure S4). We conclude that estimating JDFE parameters is in principle feasible with a modest experimental effort, at least for the purpose of ranking drug pairs.

**Figure 5.**
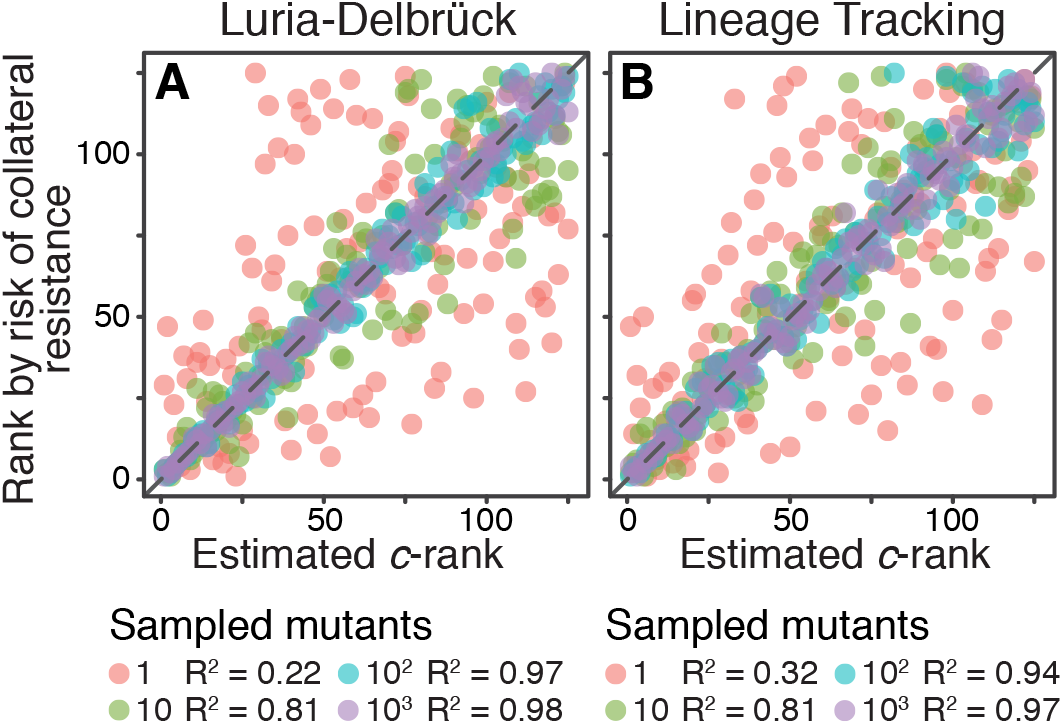
Sampling effects on the ranking of drug pairs. Both panels show correlations between the *a priori* estimated *c*-rank (*x*-axis) of the 125 Gaussian JDFEs and their *a posteriori* rank (*y*-axis) based on the observed risk of collateral resistance (same data as the *y*-axis in Figure 4C). **A.** The *c* statistic is estimated using the Luria-Delbrück method (see text for details). Cutoff for sampling mutations is 0.5*σ*, where *σ* is the standard deviation of the JDFE in the home environment. See Figure S4 for other cutoff values. **B.** The *c* statistic is estimated using the barcode lineage tracking method with *N* = 10^6^ and *U* = 10^−4^ (see text and Materials and Methods for details).

## Discussion

We have shown that many resistance mutations against multiple drugs in *E. coli* exhibit a diversity of collateral effects. If this is true more generally, it implies that there is an unavoidable uncertainty in whether any given population would evolve collateral resistance or sensitivity, which could at least in part explain inconsistencies in experimental observations. We quantified the diversity of pleiotropic effects of mutations with a joint distribution of fitness effects (JDFE) and developed a population genetic theory for predicting the expected collateral outcomes of evolution and the uncertainty around these expectations. Our theory shows that in the successional mutations regime the ensemble average rate at which fitness in the non-home environment is gained or lost during adaptation to the home environment is determined by the pleiotropy statistic *r*_2_ given by equation (8). How strongly the non-home fitness in any individual population deviates from this ensemble average is determined by the pleiotropy variance statistic *D*_22_ given by equation (11). Importantly, *r*_2_ and *D_22_* are properties of the JDFE alone, i.e., they do not depend on the parameters of any specific population. In the concurrent mutations regime, the expected rate of non-home fitness gain or loss and the associated variance are reasonably well predicted by the adjusted pleiotropy statistics 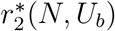 and 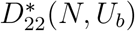. Unlike *r*_2_ and *D*_22_, the adjusted statistics depend on the population size *N* and the rate of beneficial mutations *U_b_*.

To quantitatively predict the rate or the probability of evolution of collateral drug resistance in practice would require the knowledge of both the JDFE for the focal bacterial or cancer-cell population in the presence of the specific pair of drugs and its *in vivo* population genetic parameters. Since estimating the latter parameters is very difficult, it appears unlikely that we would be able to quantitatively predict the dynamics of collateral effects, even if JDFEs were known. A more realistic application of our theory is that it allows us to rank drug pairs according to the risk of collateral resistance even when the population genetic parameters are unknown. Such robust ranking can be computed based on the collateral risk statistic 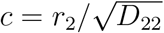, a property of the JDFE but not of the evolving population. Drug pairs with positive values of *c* have a higher chance of eliciting collateral resistance than collateral sensitivity and should be avoided; drug pairs with more negative values of *c* have a lower risk of collateral resistance evolution than those with less negative values.

What the most effective ways of measuring JDFEs are and whether it will be possible to measure JDFE *in vivo* are open questions. We speculate that the answers will depend on the shapes of the empirical JDFEs because some shapes may be more difficult to estimate than others. For example, if empirical JDFEs resemble multivariate Gaussian distributions, then we can learn all relevant parameters of such JDFE by sampling a handful of random mutants and measuring their fitness in relevant environments. One can also imagine more complex JDFEs where mutations beneficial in the home environment have a dramatically different distribution of non-home fitness effects than mutations that are deleterious or neutral in the home environment. In this case, very large samples of random mutations would be necessary to correctly predict the pleiotropic outcomes of evolution, so that methods that preferentially sample beneficial mutations may be required. We have considered two such methods, which are experimentally feasible. We have shown that both of them perform extremely well on Gaussian JDFEs in the sense that as few as 10 mutants per drug pair are sufficient to produce largely correct ranking of hypothetical drug pairs. However, it may be difficult to apply these methods *in vivo*, in which case JDFEs would have to be estimated in the lab, with selection pressures reproducing those *in vivo* as accurately as possible.

Our model relies on two important simplifications. It describes the evolution of an asexual population where all resistance alleles arise from *de novo* mutations. In reality, some resistance alleles in bacteria may be transferred horizontally (Sun et al., 2019). Understanding collateral resistance evolution in the presence of horizontal gene transfer events would require incorporating JDFE into other evolutionary dynamics models (e.g., Neher et al., 2010). Another major simplification is in the assumption that the JDFE stays constant as the population adapts. In reality the JDFE will change over time because of the depletion of the pool of adaptive mutations and because of epistasis (Good et al., 2017; Venkataram et al., 2020). How JDFEs vary among genetic backgrounds is currently unknown. In Appendix A, we have shown that our main results hold at least in the presence of a simple form of global epistasis. Empirically measuring how JDFEs vary across genotypes and theoretically understanding how such variation would affect the evolution of pleiotropic outcomes are important open question.

While we were primarily motivated by the problem of evolution of collateral drug resistance and sensitivity, our theory is applicable more broadly. The shape of JDFE must play a crucial role in determining whether the population evolves towards a generalist or diversifies into multiple specialist ecotypes. Previous literature has viewed this question primarily through the lense of two alternative hypotheses: antagonistic pleiotropy and mutation accumulation (Visher and Boots, 2020). Antagonistic pleiotropy in its strictest sense means that the population is at the Pareto front with respect to the home and non-home fitness, such that any mutation beneficial in the home environment reduces the fitness in the non-home environment (Li et al., 2019). The shape of the Pareto front then determines whether selection would favor specialists or generalists (Levins, 2020; Visher and Boots, 2020). Alternatively, a population can evolve to become a home-environment specialist even in the absence of trade-offs, simply by accumulating mutations that are neutral in the home environment but deleterious in the non-home environment (Kawecki, 1994). More recently, it has been recognized that antagonistic pleiotropy and mutation accumulation are not discrete alternatives but rather extremes of a continuum of models (Bono et al., 2020; Jerison et al., 2014, 2020). The JDFE provides a mathematical way to describe this continuum. For example, strict antagonistic pleiotropy can be modeled with a JDFE with zero probability weight in the first quadrant and a bulk of probability in the fourth quadrant. A mutation accumulation scenario can be modeled with a “+”-like JDFE where all mutations beneficial in the home environment are neutral in the non-home environment (i.e., concentrated on the *x*-axis) and all or most mutations neutral in the home environment (i.e., those on the *y*-axis) are deleterious in the non-home environment. Our theory shows that in fact all JDFEs with negative *r*_2_ lead to loss of fitness in the non-home environment and therefore can potentially promote specialization. While our theory provides this insight, further work is needed to understand how JDFEs govern adaptation to variable environments. This future theoretical work, together with empirical inquiries into the shapes of JDFEs, will not only advance our ability to predict evolution in practical situations, such as drug resistance, but it will also help us better understand the origins of ecological diversity.

## Materials and Methods

### Analysis of knock-out and deep mutational scanning data

#### Knock-out data

Chevereau et al. (2015) provide growth rate estimates for 3883 gene knock-out mutants of *E. coli* in the presence of six antibiotics. Our goal is to identify those knock-out mutations that provide resistance against one drug and are also collaterally resistant or collaterally sensitive to another drug. However, it is unclear from these original data alone which mutations have statistically significant beneficial and deleterious effects because no measurement noise estimates are provided. To address this problem, we obtained replicate wildtype growth rate measurements in the presence of antibiotics from Guillaume Chevereau and Tobias Bollenbach (available at https://github.com/ardellsarah/JDFE-project). In this additional data set, the wildtype *E. coli* strain is measured on average 476 times in the presence of each drug. We estimate the wildtype growth rate *r*_WT_ as the mean of these measurements, and we obtain the selection coefficient for all knock-out mutants as *s_i_* = *r_i_* – *r*_WT_. We also obtain the noise distribution *P*_noise_(*s*) from the replicate wildtype measurements (shown in red in the diagonal panels in Figure 1). Modeling *P*_noise_(*s*) as normal distributions, we obtain the *P*-values for each mutation in the presence of each antibiotic.

We then call any knock-out mutant as resistant against a given drug if its selection coefficient in the presence of that drug exceeds a critical value 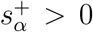. We choose 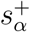 using the Benjamini-Hochberg procedure (Benjamini and Hochberg, 1995) so that the false discovery rate (FDR) among the identified resistant mutants is *α* ≈ 0.25. We could not find an 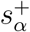 for *α* ≲ 0.25 for trimethoprim (TMP) and chloramphenicol (CHL), i.e., there were not enough knock-out mutations with positive selection coefficients to reliably distinguish them from measurement errors.

We apply the same procedure to identify mutations that are collaterally resistant and collaterally sensitive against a second drug among all mutations that are resistant against the first drug, except we aim for FDR ≲ 0.10.

#### Deep mutational scanning data

Stiffler et al. (2015) provide estimates of relative fitness for 4997 point mutations in the TEM-1 *β*-lactamase gene in the presence of cefotaxime (CEF) and four concentrations of ampicillin (AMP). They report two replicate measurements per mutant in each concentration of AMP but unfortunately only a single measurement per mutant in the presence of CEF. We chose CEF as the home environment and call all mutations with positive measured fitness effects as resistant against CEF. For each such mutation, we use two replicate measurements in each concentration of AMP to estimate its mean fitness effect and the 90% confidence interval around the mean, based on the normal distribution. We call any CEF-resistant mutation with the entire confidence interval above (below) zero as collaterally resistant (sensitive) against AMP at that concentration. All remaining CEF-resistant mutations are called collaterally neutral.

### Theory

#### Successional mutations regime

We assume that an asexual population evolves according the Wright-Fisher model in the strong selection weak mutation (SSWM) limit (Orr, 2000; Kryazhimskiy et al., 2009; Good and Desai, 2015), also known as the “successional mutations” regime (Desai and Fisher, 2007). In this regime, the population remains monomorphic until the arrival of a new mutation that is destined to fix. The waiting time for such new mutation is assumed to be much longer than the time it takes for the mutation to fix, i.e., fixation happens almost instantaneously on this time scale, after which point the population is again monomorphic. If the per genome per generation rate of beneficial mutations is *U_b_*, their typical effect is *s* and the population size is *N*, the SSWM approximation holds when *NU_b_* ≪ 1/ln (*Ns*) (Desai and Fisher, 2007).

We describe our population by a two-dimensional vector of random variables (*X_t_, Y_t_*), where *X_t_* and *Y_t_* are the population’s fitness (growth rate or the Malthusian parameter) in the home and non-home environments at generation *t*, respectively. We assume that the fitness vector of the population at the initial time point is known and is (*x*_0_, *y*_0_). We are interested in characterizing the joint probability density *p*(*x, y, t*) *dx dy* = Pr {*X_t_* ∈ [*x, x* + *dx*), *Y_t_* ∈ [*y, y* + *dy*)}.

We assume that all genotypes have the same JDFE Φ (Δ*x*, Δ*y*), i.e., there is no epistasis. In the exponential growth model, the selection coefficient of a mutation is the difference between the mutant and the ancestor growth rates in the home environment, i.e., Δ*x*. The probability of fixation of the mutant is given by Kimura’s formula, which we approximate by 2Δ*x* for Δ*x* > 0 and zero otherwise (Crow and Kimura, 1972).

If the total rate of mutations (per genome per generation) is *U*, the rate of mutations beneficial in the home environment is given by *U_b_* = U*f_b_* where 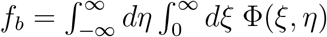 is the fraction of mutations beneficial in the home environment. Once such a mutation arises, its selection coefficients in the home and non-home environments are drawn from the JDFE of mutations beneficial in the home environment Φ_*b*_(Δ*x*, Δ*y*) = Φ(Δ*x*, Δ*y*)/*f_b_*. Then, in the SSWM limit, our population is described by a two-dimensional continuous-time continuous-space Markov chain with the transition rate from state (*x, y*) to state (*x*′, *y*′) given by

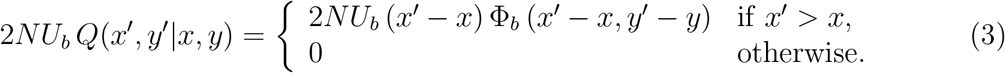

The probability distribution *p*(*x, y, t*) satisfies the integro-differential forward Kolmogorov equation (Van Kampen, 1992)

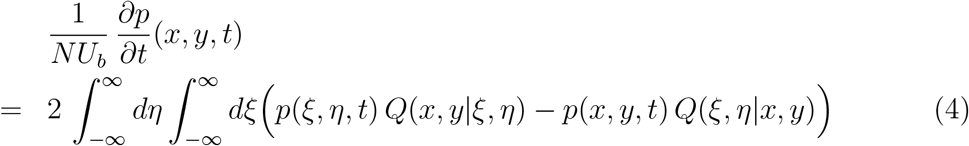

with the initial condition

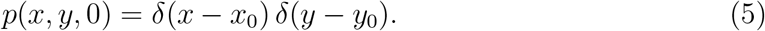

When beneficial mutations with large effects are sufficiently rare, equation (4) can be approximated by the Fokker-Planck equation (Van Kampen, 1992)

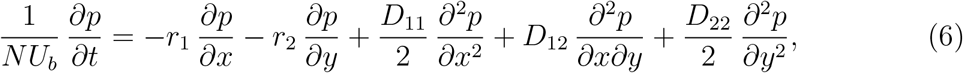

where

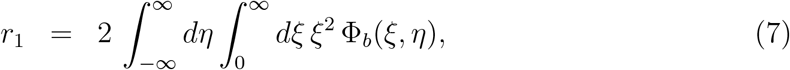

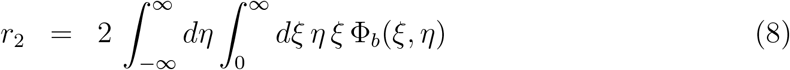

are the expected fitness effects in the home and non-home environments for a mutation fixed in the home environment, and

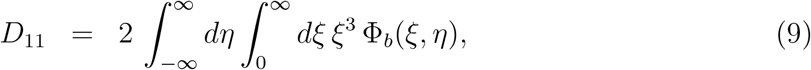

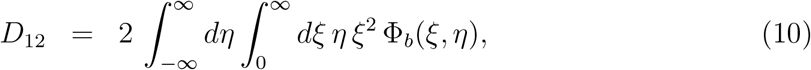

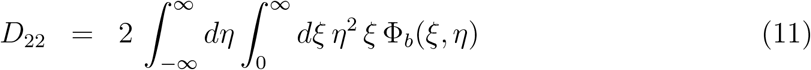

are the second moments of the distribution of the fitness effects of mutations fixed in the home environment. The solution to equation (6) with the initial condition (5) is a multi-variate normal distribution with the mean vector ***m***(*t*) and the variance-covariance matrix ***σ***^2^(*t*) given by equations (1), (2).

#### Concurrent mutations regime

The theory we developed so far for the successional mutations regime breaks down in the concurrent mutations regime, i.e., when multiple adaptive mutations segregate in the population simultaneously (Desai and Fisher, 2007). The main effect of competition between segregating adaptive lineages is that many new beneficial mutations arise in relatively low-fitness genetic backgrounds and have almost no chance of surviving competition (Desai and Fisher, 2007; Schiffels et al., 2011; Good et al., 2012). As a result, the fixation probability of a beneficial mutation with selective effect Δ*x* in the home environment is no longer 2Δ*x*. Instead, beneficial mutations that provide fitness benefits below a certain threshold *x_c_* behave as if they are effectively neutral (i.e., their fixation probability is close to zero), and most adaptation is driven by mutations with benefits above *x_c_*, where *x_c_* depends on the population genetic parameters *N* and *U_b_* as well as the shape of the distribution of fitness effects of beneficial mutations.Good et al. (2012) derived equations that allow us to calculate the effective fixation probability π*(Δ*x*; *N, U_b_*) of a beneficial mutation with the fitness benefit Δ*x* in the home environment in the concurrent mutation regime. Thus, to predict the average rate of non-home fitness change, we replace the SSWM fixation probability 2*ξ* in equation (8) with π*(*ξ*; *N*, *U_b_*) and obtain the adjusted expected pleiotropic effect

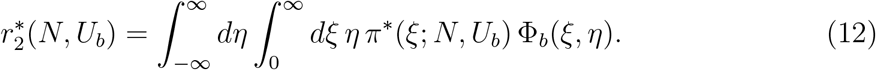

We similarly obtain the adjusted pleiotropic variance statistic

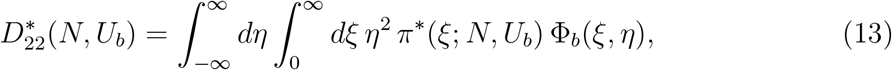

although, as discussed in Section “The population genetics of pleiotropy”, we do not expect 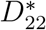 to capture all of the variation in non-home fitness trajectories.

To calculate π*(Δ*x*; *N, U_b_*) for the Gaussian JDFEs shown in Figure 2, we first substitute equation (20) in Good et al. (2012) with *β* = 2 into equations (18), (19) in Good et al. (2012) and then numerically solve these equations for *x_c_* and *v* using the Find-Root numerical method in Mathematica. Note that all our Guassian JDFEs share the same mean and variance in the home environment, so we need to solve these equations only once for each pair of *N* and *U_b_* values. We then substitute the obtained values of *x_c_* and *v* into equations (4) and (9) in Good et al. (2012) and calculate π* by a numerical integration of equation (6) in Good et al. (2012) in *R* (available at https://github.com/ardellsarah/JDFE-project).

### Ranking of drug pairs

According to equations (1), (2), both the expected non-home fitness and its variance change linearly with time, so that at time *t* the mean is 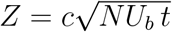 standard deviations above *y*_0_ (if *r*_2_ > 0) or below *y*_0_ (if *r*_2_ < 0), where 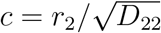. In other words, if *r*_2_ > 0, the bulk of the non-home fitness distribution eventually shifts above *y*_0_, and if *r*_2_ < 0, it shifts below *y*_0_. All else being equal, a larger value of |*c*| implies faster rate of this shift.

The interpretation of these observations in terms of collateral resistance/sensitivity is that adaptation in the presence of the first drug will eventually lead to collateral resistance against the second drug if *r*_2_ > 0 and to collateral sensitivity if *r*_2_ < 0. Furthermore, all else being equal, collateral sensitivity evolves faster and the chance of evolving collateral resistance is smaller for drug pairs with more negative *c* (i.e., larger |*c*|). Thus, we use *c* to order drug pairs from the most preferred (those with the most negative values of *c*) to least preferred (those with least negative or positive values of *c*).

### Generation of JDFEs

#### Gaussian JDFEs

The JDFEs in Figure 2 have the following parameters. Mean in the home environment: −0.05. Standard deviation in both home and non-home environments: 0.1. Means in the non-home environment: 0.08, 0.145, 0, −0.145, −0.08 in panels A through E, respectively.

The JDFEs in Figure 3 have the following parameters. Mean and standard deviation in the home environment: −0.001 and 0.01, respectively. The non-home mean varies between 0.0001 and 0.01. The non-home standard deviation varies between 0.0001 and 0.01. The correlation between home and non-home fitness varies between −0.9 and 0.9, for a total of 125 JDFEs. All parameter values and the resulting pleiotropy statistics for these JDFEs are given in the Supplementary Table S3.

#### JDFEs with equal probabilities of pleiotropically beneficial and deleterious mutations

All JDFEs in Figure S2 are mixtures of two two-dimensional uncorrelated Gaussian distributions, which have the following parameters. Mean in the home environment: 0.4. Standard deviation in both home and non-home environments: 0.1. Means in the non-home environment: 0.1 and −0.1 in panel A, 0.5 and −0.5 in panel B, 0.17 and −0.5 in panel C, and 0.5 and −0.17 in panel D.

### Simulations

We carried out two types of simulations, SSWM model simulations and full Wright-Fisher model simulations.

#### Strong selection weak mutation

The SSWM simulations were carried out using the Gillespie algorithm (Gillespie, 1976), as follows. We initiate the populations with home and non-home fitness values *x*_0_ = 0 and *y*_0_ = 0. At each iteration, we draw the waiting time until the appearance of the next beneficial mutation from the exponential distribution with the rate parameter *NU_b_* and advance the time by this amount. Then, we draw the selection coefficients Δ*x* and Δ*y* of this mutation in the home- and non-home environment, respectively, from the JDFE (a multivariate normal distribution). With probability 2Δ*x*, the mutation fixes in the population. If it does, the fitness of the population is updated accordingly.

#### Wright-Fisher model

We simulate evolution in the home environment according to the Wright-Fisher model with population size *N* as follows. We initiate the whole population with a single genotype with fitness *x*_0_ = 0 and *y*_0_ = 0 in the home and non-home environments. Suppose that at generation *t*, there are *K*(*t*) genotypes, such that genotype *i* has home- and non-home fitness *X_i_* and *Y_i_*, respectively, and it is present at frequency *f_i_*(*t*) > 0 in the population. We generate the genotype frequencies at generation *t* + 1 in three steps. In the reproduction step, we draw random numbers 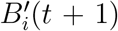, *i* = 1,…, *K*(*t*) from the multinomial distribution with the number of trials *N* and success probabilities 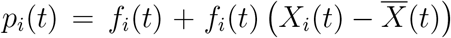, where 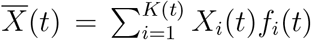 is the mean fitness of the population in the home environment at generation *t*. In the mutation step, we draw a random number *M* of new mutants from the Poisson distribution with parameter *NU*, where *U* is the total per individual per generation mutation rate. We randomly determine the “parent” genotypes in which each mutation occurs and turn the appropriate numbers of parent individuals into new mutants. We assume that each new mutation creates a new genotype and has fitness effects Δ*x* and Δ*y* in the home and non-home environments. Δ*x* and Δ*y* are drawn randomly from the JDFE Φ(Δ*x*, Δ*y*).

We obtain each mutants fitness by adding these values to the parent genotype’s home and non-home fitness values. In the final step, all genotypes that are represented by zero individuals are removed and we are left with *K*(*t* + 1) genotypes with *B_i_*(*t* + 1) > 0, *i* = 1,…, *K*(*t* + 1) individuals. Then we set *f_i_*(*t* + 1) = *B_i_*(*t* + 1)/*N*.

### Sampling beneficial mutants from JDFEs and estimating the *c* statistic

We model the LD sampling method by randomly drawing mutants from the JDFE until the desired number *K* of mutants whose home fitness exceeds the focal threshold are sampled. We estimate the *c* statistic from the pairs of home and non-home fitness effects *X_i_* and *Y_i_* of these *i* = 1,…, *K* sampled mutants. To do so, we first estimate *r*_2_ and *D*_22_ as 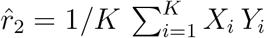 and 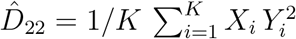. We then calculate 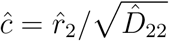.

For the BLT sampling method, we simulate the Wright-Fisher model as described above for *N* = 10^6^ and *U* = 10^−4^ for 250 generations. At generation 250, we randomly sample existing beneficial mutants proportional to their frequency in the population without replacement (i.e., the same beneficial mutation is sampled at most once). Sampling more than ~ 50 distinct beneficial mutants from a single population becomes difficult because there may simply be not enough such mutants or some of them may be at very low frequencies. Therefore, if the desired number of mutants to sample exceeds 50, we run multiple replicate simulations and sample a maximum of 100 distinct beneficial mutants per replicate until the desired number of mutants is reached. We then estimate the c statistics as with the LD method.

## Code availability

All scripts are available at https://github.com/ardellsarah/JDFE-project.

## Acknowledgements

We thank Shea Summers and Flora Tang for help and input at the initial stages of the project and Kryazhimskiy and Meyer labs for feedback. We thank Tobias Bollenbach and Guillaume Chevereau for providing wildtype growth rate measurement data. This work was supported by the BWF Career Award at Scientific Interface (Grant 1010719.01), the Alfred P. Sloan Foundation (Grant FG-2017-9227), the Hellman Foundation and NIH (Grant 1R01GM137112).

## Supplementary Figures

**Supplementary Figure S1.**
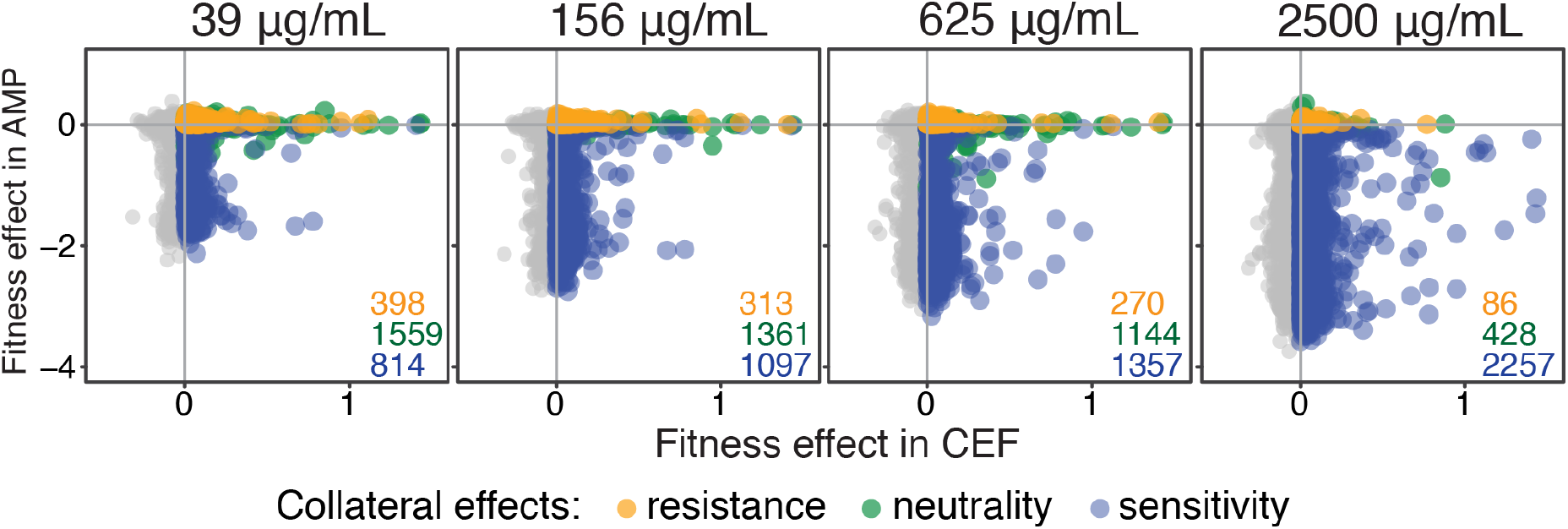
Fitness effects of single point mutations in the TEM-1 *β*-lactamase gene in *E. coli* in the presence of cefotaxime and ampicillin. Data from Stiffler et al. (2015). Panels show data for different concentrations of ampicillin, as indicated. Fitness is measured as the change in the log ratio of the mutant to wildtype frequency during growth in the presence of the drug. Cefotaxime (CEF) is chosen as the home environment (see Materials and Methods for details). Each point represents a single point mutation and is colored by its (collateral) fitness effect in the presence of ampicillin, as indicated in the legend. The numbers of mutations with positive fitness in the presence of cefotaxime with different collateral effects are shown in the lower right corner of each panel.

**Supplementary Figure S2.**
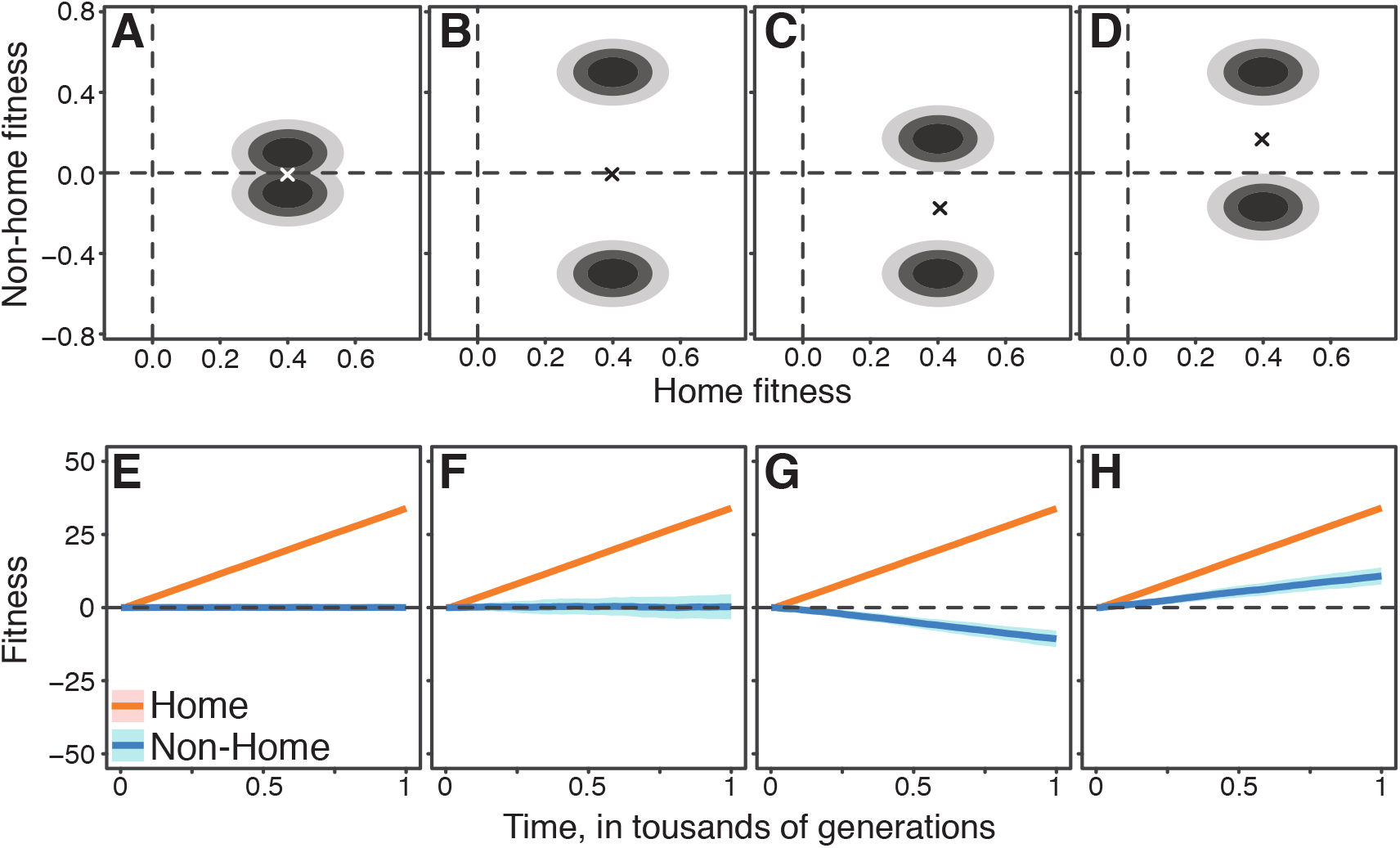
Same as Figure 2, but for JDFEs with equal probability weights in the first and fourth quadrants. See Materials and Methods for details.

**Supplementary Figure S3.**
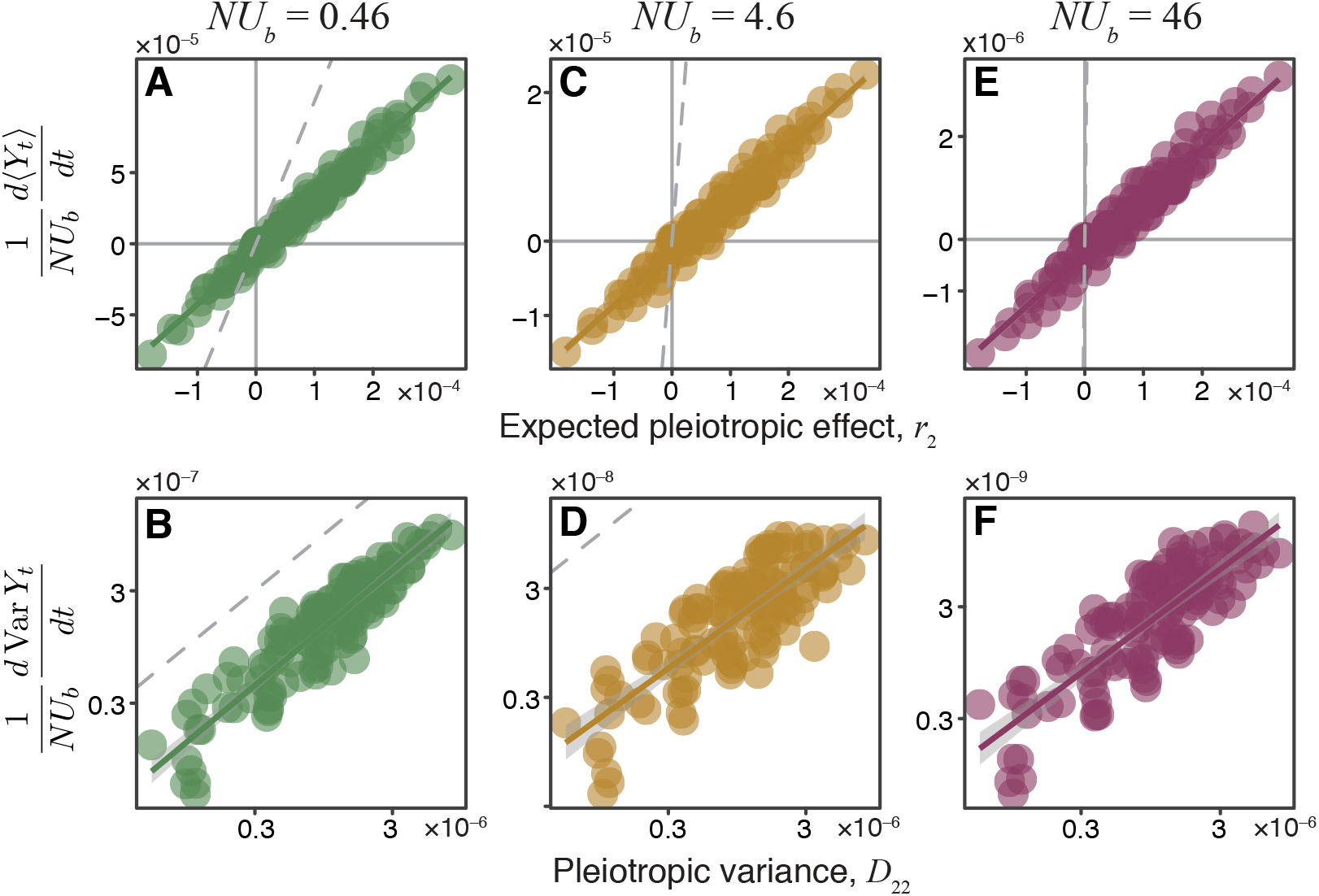
Same as Figure 3C–H, but with *r*_2_ and *D*_22_ shown on the *x*-axis.

**Supplementary Figure S4.**
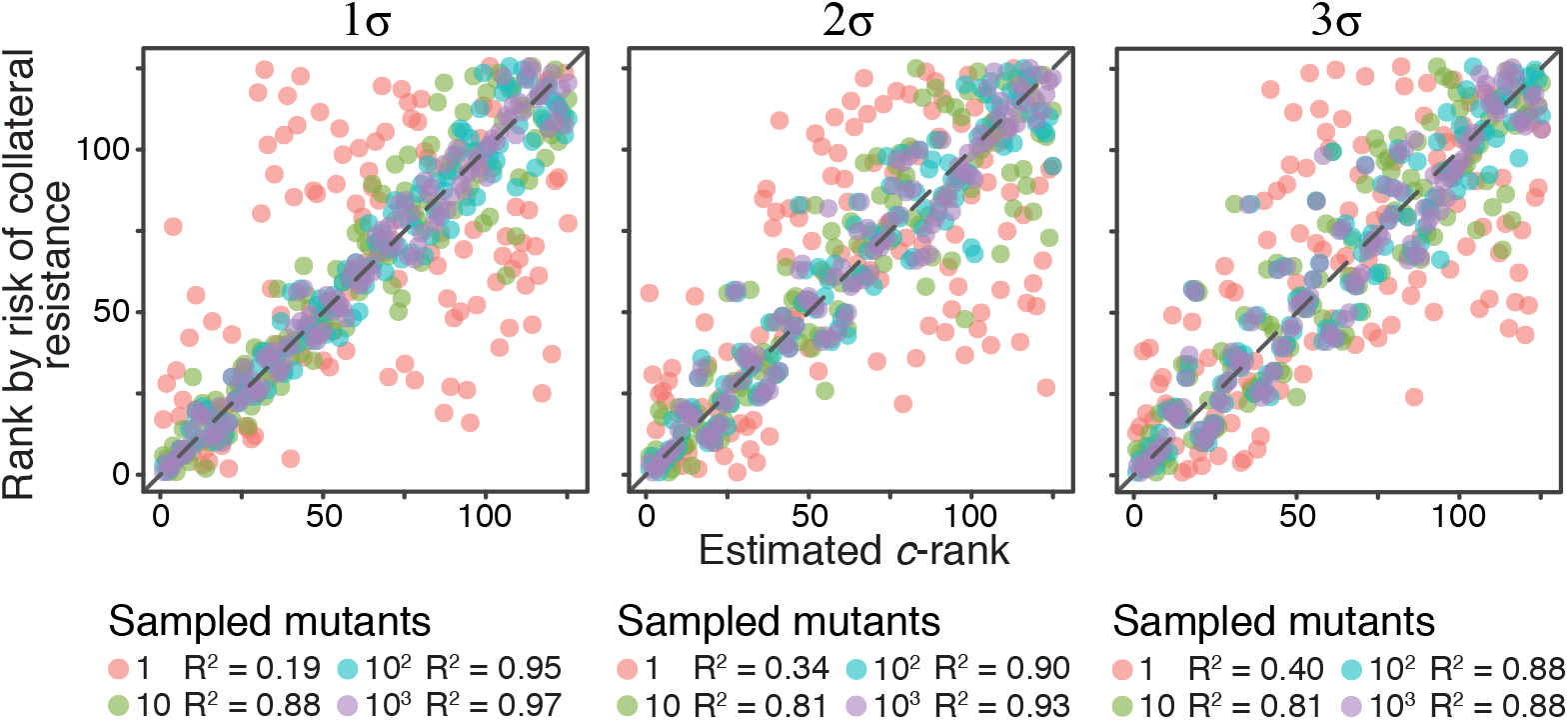
Same as Figure 5A, but with different thresholds for sampling mutations, as indicated above each panel (*σ* is the standard deviation of the JDFE in the home environment). See Materials and Methods for details.

## Supplementary Tables

**Supplementary Table S1.** *P*-values and calls of collateral effects of beneficial knock-out mutations in the Chevereau et al. (2015) data (see Materials and Methods for details).

**Supplementary Table S2.** Calls of collateral effects of mutations beneficial in CEF in the Stiffler et al. (2015) data (see Materials and Methods for details).

**Supplementary Table S3.** Parameters and summary statistics of simulation results for all Gaussian JDFEs used in Figure 3.

## Appendix A JDFE with global epistasis

Results in the main text were derived under the assumption that all genotypes have the same JDFE, i.e., in the absence of epistasis. In reality, JDFEs probably vary from one genotype to another, but how they vary is not yet well characterized. Recent studies have found that the fitness effects of many mutations available to a genotype in a given environment depend primarily on the fitness of that genotype in that environment (Khan et al., 2011; Chou et al., 2011; Wiser et al., 2013; Kryazhimskiy et al., 2014; Johnson et al., 2019; Wang et al., 2016; Aggeli et al., 2020; Lukačišinová et al., 2020). This dependence is sometimes referred to as global or fitness-dependent epistasis (Kryazhimskiy et al., 2009, 2014; Reddy and Desai, 2020; Husain and Murugan, 2020). Here, we ask whether our main results would hold if the pathogen population evolves on a JDFE with global epistasis.

Global epistasis can be modeled in our framework by assuming that the JDFE Φ*_g_* of genotype *g* depends only the fitness of this genotype in the home and non-home environments, *x*(*g*), *y*(*g*), i.e. Φ_*g*_ (Δ*x*, Δ*y*) = Φ_*x*(*g*),*y*(*g*)_ (Δ*x*, Δ*y*), which is a two-dimensional extension of the model considered by Kryazhimskiy et al. (2009). Thus, in the SSWM regime, the population can still be fully described by its current pair of fitness values in the home and non-home environments (*X_t_, Y_t_*). The dynamics of the probability density *p*(*x, y, t*) are governed by the same Kolmogorov equation as in the non-epistatic case, which can still be approximated by a diffusion equation (6). However, while in the non-epistatic case the drift and diffusion coefficients of this equation, *r*_2_, *r*_2_, *D*_11_, *D*_12_ and *D*_22_ are constants, in the presence of global epistasis, they become functions of *x* and *y*. Although this equation cannot be solved analytically in the general case, it can be solved numerically, provided that the functions *r*_1_(*x,y*), *r*_2_(*x,y*), *D*_11_(*x,y*), *D*_12_(*x,y*) and *D*_22_(*x,y*) are known. Thus, in principle, our theory can predict the trajectories of non-home fitness in the presence of global epistasis.

To explore the implications of global epistasis for collateral drug resistance evolution, we consider the simplest scenario where the functional form of global epistasis (i.e., how Φ*_x,y_* depends on *x* and *y*) is the same across different drugs. In this case, we would expect that the ranking of drug pairs according to the risk of collateral resistance would be the same for all genotypes. In particular, the drug pair whose risk of collateral resistance risk is the lowest for the wildtype should also be the pair with the lowest risk for the evolved genotypes.

To test this prediction, we model resistance evolution on Gaussian JDFEs whose mean vector and the correlation coefficient are fixed while the standard deviations *σ*_h_(*x*) and *σ*_nh_(*y*) in the home and non-home environments decrease linearly with the fitness in the respective environment, *σ*_h_(*x*) = max {0, *σ*_h,0_ – *γ*_h_ *x*} and *σ*_nh_(*y*) = max {0, *σ*_nh,0_ – *γ*_nh_ *y*}. Appendix 1 Figure 1A shows how one such JDFE changes along an expected evolutionary trajectory. The corresponding expected home and non-home fitness trajectories and their variance are shown in Appendix 1 Figure 1B. Appendix 1 Figure 1C shows how the probability (risk) of collateral resistance changes over time on four different JDFEs with global epistasis. For the ancestral strain (whose fitness we set by convention to *x* = *y* = 0), these four JDFEs are identical to those shown in Figure 4A; as the populations evolve, JDFEs change as specified above with *γ*_h_ = *γ* _nh_ = 0.5. As expected, the ranking of these epistatic JDFEs according to the risk of collateral resistance stays constant over time and can be predicted from estimates of the *c* parameters for the ancestral strain.

**Appendix 1 Figure 1.**
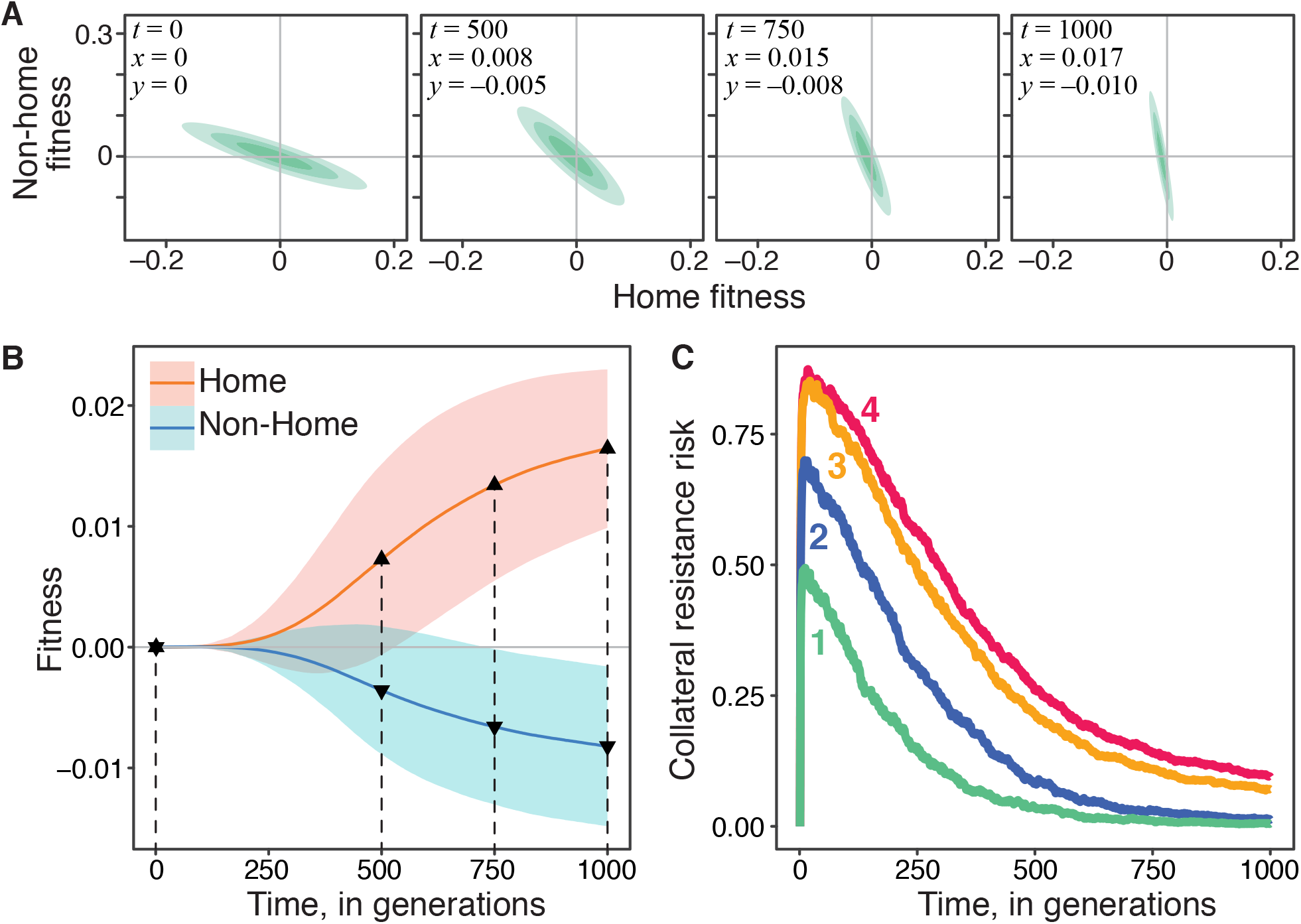
Evolution on JDFEs with global epistasis and the risk of collateral resistance. **A.** Gaussian JDFE with global epistasis as it changes along the expected evolutionary trajectory shown in panel B. Parameters of the initial JDFE at *x* = *y* = 0 are the same as for the rank 1 JDFE in Figure 4A; *γ*_h_ = *γ*_nh_ = 0.5. **B.** Home and non-home fitness trajectories for the JDFE with global epistasis shown in panel A. Thick lines show the mean, ribbons show ±1 standard deviation estimated from 500 replicate simulations. Population size *N* = 10^4^, mutation rate *U* = 10^−4^. Dashed vertical lines indicate the time points at which the JDFE snapshots in panel A are shown. **C.** Probability of collateral resistance over time for four Gaussian JDFE with global epistasis. Parameters of the initial JDFEs at *x* = *y* = 0 are the same as for the four JDFE in Figure 4A, and *γ*_h_ = *γ*_nh_ = 0.5 for all of them. *N* = 10^4^, mutation rate *U* = 10^−4^, 1500 replicate simulation runs per JDFE. Colored numbers indicate the predicted *c*-rank of the initial JDFEs (same as in Figure 4A).

## References

Aggeli, D., Y. Li and G. Sherlock. 2020. Changes in the distribution of fitness effects and adaptive mutational spectra following a single first step towards adaptation. bioRxiv: 2020.06.12.148833 URL https://doi.org/10.1101/2020.06.12.148833.

Andersson, D. I. and D. Hughes. 2014. Microbiological effects of sublethal levels of antibiotics. Nature Reviews Microbiology 12:465–478.

Bakerlee, C. W., A. M. Phillips, A. N. N. Ba and M. M. Desai. 2021. Dynamics and variability in the pleiotropic effects of adaptation in laboratory budding yeast populations. bioRxiv.

Barbosa, C., V. Trebosc, C. Kemmer, P. Rosenstiel, R. Beardmore, H. Schulenburg and G. Jansen. 2017. Alternative evolutionary paths to bacterial antibiotic resistance cause distinct collateral effects. Molecular biology and evolution 34:2229–2244.

Barton, N. 1990. Pleiotropic models of quantitative variation. Genetics 124:773–782.

Batra, A., R. Roemhild, E. Rousseau, S. Franzenburg, S. Niemann and H. Schulenburg. 2021. High potency of sequential therapy with only beta-lactam antibiotics. Elife 10:e68876.

Benjamini, Y. and Y. Hochberg. 1995. Controlling the false discovery rate: a practical and powerful approach to multiple testing. Journal of the Royal statistical society: series B (Methodological) 57:289–300.

Bergstrom, C. T., M. Lo and M. Lipsitch. 2004. Ecological theory suggests that antimicrobial cycling will not reduce antimicrobial resistance in hospitals. Proceedings of the National Academy of Sciences 101:13285–13290.

Blundell, J. R., K. Schwartz, D. Francois, D. S. Fisher, G. Sherlock and S. F. Levy. 2019. The dynamics of adaptive genetic diversity during the early stages of clonal evolution. Nature ecology & evolution 3:293–301.

Bono, L. M., J. A. Draghi and P. E. Turner. 2020. Evolvability costs of niche expansion. Trends in Genetics 36:14–23.

Bono, L. M., L. B. Smith Jr, D. W. Pfennig and C. L. Burch. 2017. The emergence of performance trade-offs during local adaptation: insights from experimental evolution. Molecular ecology 26:1720–1733.

Card, K. J., J. A. Jordan and R. E. Lenski. 2021. Idiosyncratic variation in the fitness costs of tetracycline-resistance mutations in escherichia coli. Evolution 75:1230–1238.

Card, K. J., M. D. Thomas, J. L. Graves Jr, J. E. Barrick and R. E. Lenski. 2020. Genomic evolution of antibiotic resistance is contingent on genetic background following a long-term experiment with *Escherichia coli*. bioRxiv: 2020.08.19.258384 URL https://doi.org/10.1101/2020.08.19.258384.

Chevereau, G., M. Dravecká, T. Batur, A. Guvenek, D. H. Ayhan, E. Toprak and T. Bollenbach. 2015. Quantifying the determinants of evolutionary dynamics leading to drug resistance. PLoS Biology 13:e1002299.

Chou, H.-H., H.-C. Chiu, N. F. Delaney, D. Segrè and C. J. Marx. 2011. Diminishing returns epistasis among beneficial mutations decelerates adaptation. Science 332:1190–1192.

Connallon, T. and A. G. Clark. 2015. The distribution of fitness effects in an uncertain world. Evolution 69:1610–1618.

Crow, J. F. and M. Kimura. 1972. An introduction to population genetics theory. Harper & Row Ltd.

Das, S. G., S. O. Direito, B. Waclaw, R. J. Allen and J. Krug. 2020. Predictable properties of fitness landscapes induced by adaptational tradeoffs. eLife 9:e55155.

Desai, M. M. and D. S. Fisher. 2007. Beneficial mutation–selection balance and the effect of linkage on positive selection. Genetics 176:1759–1798.

Eyre-Walker, A. and P. D. Keightley. 2007. The distribution of fitness effects of new mutations. Nature Reviews Genetics 8:610–618.

Gillespie, D. T. 1976. A general method for numerically simulating the stochastic time evolution of coupled chemical reactions. Journal of computational physics 22:403–434.

Gjini, E. and K. B. Wood. 2021. Price equation captures the role of drug interactions and collateral effects in the evolution of multidrug resistance. Elife 10:e64851.

Good, B. H. and M. M. Desai. 2015. The impact of macroscopic epistasis on long-term evolutionary dynamics. Genetics 199:177–190.

Good, B. H., M. J. McDonald, J. E. Barrick, R. E. Lenski and M. M. Desai. 2017. The dynamics of molecular evolution over 60,000 generations. Nature 551:45–50.

Good, B. H., I. M. Rouzine, D. J. Balick, O. Hallatschek and M. M. Desai. 2012. Distribution of fixed beneficial mutations and the rate of adaptation in asexual populations. Proceedings of the National Academy of Sciences 109:4950–4955.

Harmand, N., R. Gallet, R. Jabbour-Zahab, G. Martin and T. Lenormand. 2017. Fisher’s geometrical model and the mutational patterns of antibiotic resistance across dose gradients. Evolution 71:23–37.

Hietpas, R. T., C. Bank, J. D. Jensen and D. N. Bolon. 2013. Shifting fitness landscapes in response to altered environments. Evolution 67:3512–3522.

Husain, K. and A. Murugan. 2020. Physical constraints on epistasis. Molecular Biology and Evolution URL https://doi.org/10.1093/molbev/msaa124. In press.

Imamovic, L. and M. O. Sommer. 2013. Use of collateral sensitivity networks to design drug cycling protocols that avoid resistance development. Science translational medicine 5:204ra132.

Jensen, P., B. Holm, M. Sorensen, I. Christensen and M. Sehested. 1997. In vitro cross-resistance and collateral sensitivity in seven resistant small-cell lung cancer cell lines: preclinical identification of suitable drug partners to taxotere, taxol, topotecan and gemcitabin. British journal of cancer 75:869–877.

Jerison, E. R., S. Kryazhimskiy and M. M. Desai. 2014. Pleiotropic consequences of adaptation across gradations of environmental stress in budding yeast. arXiv:1409.7839 URL https://arxiv.org/abs/1409.7839.

Jerison, E. R., A. N. Nguyen Ba, M. M. Desai and S. Kryazhimskiy. 2020. Chance and necessity in the pleiotropic consequences of adaptation for budding yeast. Nature Ecology & Evolution 4:601–611.

Johnson, M. S., A. Martsul, S. Kryazhimskiy and M. M. Desai. 2019. Higher-fitness yeast genotypes are less robust to deleterious mutations. Science 366:490–493.

Johnson, T. and N. Barton. 2005. Theoretical models of selection and mutation on quantitative traits. Philosophical Transactions of the Royal Society B: Biological Sciences 360:1411–1425.

Jones, A. G., S. J. Arnold and R. Bürger. 2003. Stability of the g-matrix in a population experiencing pleiotropic mutation, stabilizing selection, and genetic drift. Evolution 57:1747–1760.

Kawecki, T. J. 1994. Accumulation of deleterious mutations and the evolutionary cost of being a generalist. The American Naturalist 144:833–838.

Khan, A. I., D. M. Dinh, D. Schneider, R. E. Lenski and T. F. Cooper. 2011. Negative epistasis between beneficial mutations in an evolving bacterial population. Science 332:1193–1196.

Kimura, M. 1962. On the probability of fixation of mutant genes in a population. Genetics 47:713.

King, J. L. 1972. The role of mutation in evolution. In Proceedings of the Sixth Berkeley Symposium on Mathematical Statistics and Probability, vol. 5, 69–100. University of California Press Berkeley.

Kryazhimskiy, S., D. P. Rice, E. R. Jerison and M. M. Desai. 2014. Global epistasis makes adaptation predictable despite sequence-level stochasticity. Science 344:1519–1522.

Kryazhimskiy, S., G. Tkačik and J. B. Plotkin. 2009. The dynamics of adaptation on correlated fitness landscapes. Proceedings of the National Academy of Sciences 106:18638–18643.

Lande, R. and S. J. Arnold. 1983. The measurement of selection on correlated characters. Evolution 37:1210–1226.

Lang, G. I., D. P. Rice, M. J. Hickman, E. Sodergren, G. M. Weinstock, D. Botstein and M. M. Desai. 2013. Pervasive genetic hitchhiking and clonal interference in forty evolving yeast populations. Nature 500:571–574.

Lázár, V., A. Martins, R. Spohn, L. Daruka, G. Grézal, G. Fekete, M. Számel, P. K. Jangir, B. Kintses, B. Csörgő et al. 2018. Antibiotic-resistant bacteria show widespread collateral sensitivity to antimicrobial peptides. Nature microbiology 3:718–731.

Levins, R. 2020. Evolution in changing environments. Princeton University Press.

Levy, S. F., J. R. Blundell, S. Venkataram, D. A. Petrov, D. S. Fisher and G. Sherlock. 2015. Quantitative evolutionary dynamics using high-resolution lineage tracking. Nature 519:181–186.

Li, Y., D. A. Petrov and G. Sherlock. 2019. Single nucleotide mapping of trait space reveals pareto fronts that constrain adaptation. Nature ecology & evolution 3:1539–51.

Lindsey, H. A., J. Gallie, S. Taylor and B. Kerr. 2013. Evolutionary rescue from extinction is contingent on a lower rate of environmental change. Nature 494:463–467.

Lukačišinová, M., B. Fernando and T. Bollenbach. 2020. Highly parallel lab evolution reveals that epistasis can curb the evolution of antibiotic resistance. Nature communications 11:1–14.

MacLean, R. C. and A. Buckling. 2009. The distribution of fitness effects of beneficial mutations in *Pseudomonas aeruginosa*. PLoS Genet 5:e1000406.

Maltas, J., D. M. McNally and K. B. Wood. 2019. Evolution in alternating environments with tunable inter-landscape correlations. bioRxiv: 803619 URL https://doi.org/10.1101/803619.

Maltas, J. and K. B. Wood. 2019. Pervasive and diverse collateral sensitivity profiles inform optimal strategies to limit antibiotic resistance. PLoS Biology 17:e3000515.

Martin, G. and T. Lenormand. 2008. The distribution of beneficial and fixed mutation fitness effects close to an optimum. Genetics 179:907–916.

Martin, G. and T. Lenormand. 2015. The fitness effect of mutations across environments: Fisher’s geometrical model with multiple optima. Evolution 69:1433–1447.

Mira, P. M., J. C. Meza, A. Nandipati and M. Barlow. 2015. Adaptive landscapes of resistance genes change as antibiotic concentrations change. Molecular biology and evolution 32:2707–2715.

Neher, R. A., B. I. Shraiman and D. S. Fisher. 2010. Rate of adaptation in large sexual populations. Genetics 184:467–481.

Nguyen Ba, A. N., I. Cvijović, J. I. R. Echenique, K. R. Lawrence, A. Rego-Costa, X. Liu, S. F. Levy and M. M. Desai. 2019. High-resolution lineage tracking reveals travelling wave of adaptation in laboratory yeast. Nature 575:494–499.

Nichol, D., J. Rutter, C. Bryant, A. M. Hujer, S. Lek, M. D. Adams, P. Jeavons, A. R. Anderson, R. A. Bonomo and J. G. Scott. 2019. Antibiotic collateral sensitivity is contingent on the repeatability of evolution. Nature communications 10:1–10.

Ohta, T. 1987. Very slightly deleterious mutations and the molecular clock. Journal of Molecular Evolution 26:1–6.

Orr, H. A. 2000. The rate of adaptation in asexuals. Genetics 155:961–968.

Orr, H. A. 2003. The distribution of fitness effects among beneficial mutations. Genetics 163:1519–1526.

Oz, T., A. Guvenek, S. Yildiz, E. Karaboga, Y. T. Tamer, N. Mumcuyan, V. B. Ozan, G. H. Senturk, M. Cokol, P. Yeh et al. 2014. Strength of selection pressure is an important parameter contributing to the complexity of antibiotic resistance evolution. Molecular biology and evolution 31:2387–2401.

Paaby, A. B. and M. V. Rockman. 2013. The many faces of pleiotropy. Trends in genetics 29:66–73.

Pál, C., B. Papp and V. Lázár. 2015. Collateral sensitivity of antibiotic-resistant microbes. Trends in microbiology 23:401–407.

Pinheiro, F., O. Warsi, D. I. Andersson and M. Läassig. 2021. Metabolic fitness landscapes predict the evolution of antibiotic resistance. Nature Ecology & Evolution 1–11.

Pluchino, K. M., M. D. Hall, A. S. Goldsborough, R. Callaghan and M. M. Gottesman. 2012. Collateral sensitivity as a strategy against cancer multidrug resistance. Drug Resistance Updates 15:98–105.

Qian, W., D. Ma, C. Xiao, Z. Wang and J. Zhang. 2012. The genomic landscape and evolutionary resolution of antagonistic pleiotropy in yeast. Cell reports 2:1399–1410.

Reddy, G. and M. M. Desai. 2020. Global epistasis emerges from a generic model of a complex trait. bioRxiv: 2020.06.14.150946 URL https://www.biorxiv.org/content/early/2020/06/14/2020.06.14.150946.

Rees, K. and T. Bataillon. 2006. Distribution of fitness effects among beneficial mutations before selection in experimental populations of bacteria. Nature Genetics 38:484–488.

Roemhild, R., M. Linkevicius and D. I. Andersson. 2020. Molecular mechanisms of collateral sensitivity to the antibiotic nitrofurantoin. PLoS Biology 18:e3000612.

Roff, D. A. and D. Fairbairn. 2007. The evolution of trade-offs: where are we? Journal of evolutionary biology 20:433–447.

Rose, M. R. 1982. Antagonistic pleiotropy, dominance, and genetic variation. Heredity 48:63–78.

Schiffels, S., G. J. Szöllősi, V. Mustonen and M. Lässig. 2011. Emergent neutrality in adaptive asexual evolution. Genetics 189:1361–1375.

Shoval, O., H. Sheftel, G. Shinar, Y. Hart, O. Ramote, A. Mayo, E. Dekel, K. Kavanagh and U. Alon. 2012. Evolutionary trade-offs, Pareto optimality, and the geometry of phenotype space. Science 336:1157–1160.

Slatkin, M. and S. A. Frank. 1990. The quantitative genetic consequences of pleiotropy under stabilizing and directional selection. Genetics 125:207–213.

Stiffler, M. A., D. R. Hekstra and R. Ranganathan. 2015. Evolvability as a function of purifying selection in TEM-1 *β*-lactamase. Cell 160:882–892.

Sun, D., K. Jeannot, Y. Xiao and C. W. Knapp. 2019. Horizontal gene transfer mediated bacterial antibiotic resistance. Frontiers in microbiology 10:1933.

Tikhonov, M., S. Kachru and D. S. Fisher. 2020. A model for the interplay between plastic tradeoffs and evolution in changing environments. Proceedings of the National Academy of Sciences 117:8934–8940.

Van Kampen, N. G. 1992. Stochastic processes in physics and chemistry, vol. 1. Elsevier.

Van Opijnen, T., K. L. Bodi and A. Camilli. 2009. Tn-seq: high-throughput parallel sequencing for fitness and genetic interaction studies in microorganisms. Nature methods 6:767–772.

Venkataram, S., B. Dunn, Y. Li, A. Agarwala, J. Chang, E. R. Ebel, K. Geiler-Samerotte, L. Hérissant, J. R. Blundell, S. F. Levy et al. 2016. Development of a comprehensive genotype-to-fitness map of adaptation-driving mutations in yeast. Cell 166:1585–1596.

Venkataram, S., R. Monasky, S. H. Sikaroodi, S. Kryazhimskiy and B. Kaçar. 2020. Evolutionary stalling and a limit on the power of natural selection to improve a cellular module. Proceedings of the National Academy of Sciences 117:18582–18590.

Visher, E. and M. Boots. 2020. The problem of mediocre generalists: population genetics and eco-evolutionary perspectives on host breadth evolution in pathogens. Proceedings of the Royal Society B 287:20201230.

Wagner, G. P. and J. Zhang. 2011. The pleiotropic structure of the genotype–phenotype map: the evolvability of complex organisms. Nature Reviews Genetics 12:204–213.

Wang, S. and L. Dai. 2019. Evolving generalists in switching rugged landscapes. PLoS Comput Biology 15:e1007320.

Wang, Y., C. D. Arenas, D. M. Stoebel, K. Flynn, E. Knapp, M. M. Dillon, A. Wünsche, P. J. Hatcher, F. B.-G. Moore, V. S. Cooper et al. 2016. Benefit of transferred mutations is better predicted by the fitness of recipients than by their ecological or genetic relatedness. Proceedings of the National Academy of Sciences 113:5047–5052.

Wiser, M. J., N. Ribeck and R. E. Lenski. 2013. Long-term dynamics of adaptation in asexual populations. Science 342:1364–1367.

